# TLPath predicts telomere length in human tissues from histopathology images

**DOI:** 10.1101/2025.03.04.641489

**Authors:** Anamika Yadav, Kyle Alvarez, Akanimoh Adeleye, Yu Xin Wang, Sanju Sinha

## Abstract

Telomere dysfunction is a key hallmark of aging linked to numerous age-related diseases including cardiovascular disorders, pulmonary fibrosis, and metabolic syndromes. Despite decades of research yielding strong evidence linking telomere biology to aging processes, the field faces a critical bottleneck: current telomere measurement methods require specialized molecular techniques that prevent large-scale studies and clinical implementation. Here we present TLPath, a novel deep learning framework that extracts normal tissue architecture from routine histopathology (H&E) images to predict bulk-tissue telomere length. Trained on the Genotype-Tissue Expression cohort comprising >7.3 million patch images from >5,000 whole-slide images across 919 individuals, TLPath makes a remarkable discovery: the extracted morphological features spontaneously separate young, middle-aged, and elderly individuals within most tissue types— demonstrating for the first time that aging causes substantial architectural changes in tissues detectable without explicit age supervision. These extracted features can predict bulk-telomere length with significant accuracy (>0.51 in well-represented tissues), outperforming chronological age as a predictor (correlation = 0.20) and identifying age-discordant cases – detecting both accelerated telomeres shortening in young individuals and preserved telomeres in older individuals. Mechanistic interpretation reveals that TLPath leverages established senescence morphological markers, including nuclear-to-cytoplasmic ratio and nuclear shape variation, for its predictions. We applied TLPath in ∼2,800 new GTEx biopsies where concordant with known association, the predicted telomere length is shorter across most tissues from individuals with Type 1/2 diabetes. Overall, we demonstrate that aging substantially alters tissue morphology, which TLPath captures and uses to predict telomere length, enabling large-scale telomere biology studies using existing tissue archives.

## 1. INTRODUCTION

Telomeres, a nucleoprotein complex at chromosome ends, serve as guardians of genomic integrity by preventing chromosome ends from being recognized as DNA double-strand breaks (1). These complexes progressively shorten with each cell division, eventually compromising their protective ability. This triggers a DNA damage response leading to cellular senescence (2). Importantly, even a few critically short telomeres are sufficient to trigger senescence regardless of average telomere length (2). Beyond length-dependent dysfunction, oxidative damage to telomere can also cause its dysfunction without shortening (3), highlighting that telomere biology extends beyond just length measurements. Overall, telomere dysfunction has emerged as a key driver of multiple age-related pathologies, with blood-based telomere length shown to be a predictor of disease progression and mortality in conditions ranging from idiopathic pulmonary fibrosis to cardiovascular diseases (2, 4). Thus, telomere shortening has been recognized as one of the aging hallmarks (5).

Considering its importance, multiple methods have evolved to measure telomere length, each with distinct trade-offs (2). Southern blotting measures bulk-tissue (mean across cell population in the biopsy) telomere length using terminal restriction fragments (6), while quantitative PCR determines the ratio of telomere repeat copy number to single copy gene number (7). More specialized techniques like single telomere length analysis (STELA) (8) and telomere shortest length assay (TeSLA) can measure individual telomeres, including the shortest ones. Flow-FISH and qFISH enable single-cell resolution through fluorescence in situ hybridization with telomere peptide nucleic acid probes, though they cannot detect telomeres below a threshold sufficient for probe hybridization (9). Recently, a non-PCR Luminex platform has enabled large-scale bulk measurements across tissues, a critical resource of 24 human tissues revealing tissue-specific variations and age-dependent shortening in a few tissues (10). However, all current approaches require either tissue homogenization or complex molecular techniques, creating a critical barrier for large-scale studies of telomere dynamics in intact tissue (2).

Prior studies have identified easily quantifiable cellular morphological changes associated with telomere shortening, primarily through its role in triggering cellular senescence (2). Cells with short telomeres display characteristic changes including increased size, irregular shape, enhanced cytoplasmic granularity, and formation of senescence-associated heterochromatin foci (SAHF) (11). Recent advances in computational pathology have demonstrated that high-resolution tissue images can predict various molecular properties, including mutation status, gene expression profiles, and chromosomal alterations. This raises the possibility that cellular morphology from routinely available tissue histology might have predictive power to determine tissue telomere length, potentially enabling scalable measurement without the need for specialized molecular techniques.

To address this need, we developed TLPath, a computational framework that predicts bulk-tissue telomere length directly from routine histopathology (H&E) images. We hypothesized that by systematically analyzing cellular morphology in whole-slide images, we could develop a model to predict bulk telomere length measurements. To test this, we leveraged the Genotype-Tissue Expression (GTEx) project’s unique resource of paired H&E slides and Luminex-based telomere length measurements (10), comprising 7,304,323 patch images from 5,263 whole-slide images across 919 individuals spanning 18 tissue types. We recognize that the shortest telomeres and telomere dysfunction independent of length are emerging as more precise measures of telomere biology, but we focus on bulk measurements due to the availability of large-scale training data for model development. TLPath enables for the first time the prediction of bulk-tissue telomere length directly from standard histopathology images, potentially transforming our ability to study telomere biology at scale.

## 2. RESULTS

### 2.1. Overview of TLPath

TLPath is a computational framework designed to predict bulk-tissue telomere length (henceforth called just *telomere length*) in normal tissues from H&E-stained histopathology images. <u>It is based on the hypothesis, hinted in prior studies (12, 13), that cell and tissue morphology visible in tissue H&E-image can accurately determine the bulk-telomere length of the tissue</u>. Accordingly, the input of TLPath is an H&E-stained slide image of normal human tissue and the output is the predicted length of tissue telomere. TLPath comprises of four sequential steps (**Figure 1a**) 1) starting with slide preprocessing, 2) followed by morphological features extraction, 3) then creating a whole slide image (WSI)-representation, and, finally, 4) building a supervised model to predict tissue telomere length based on morphological features (details below and in **Methods 4.3**). TLPath is trained on Genotype-Tissue Expression (GTEx) project cohort (**Figure 2a**), a public cohort of tissue samples from postmortem donors. Here, relative telomere length (RTL) is determined by quantifying the telomere repeat abundance in a DNA sample relative to a standard reference sample for each tissue and validated using southern blot and qPCR.

**Figure 1:**
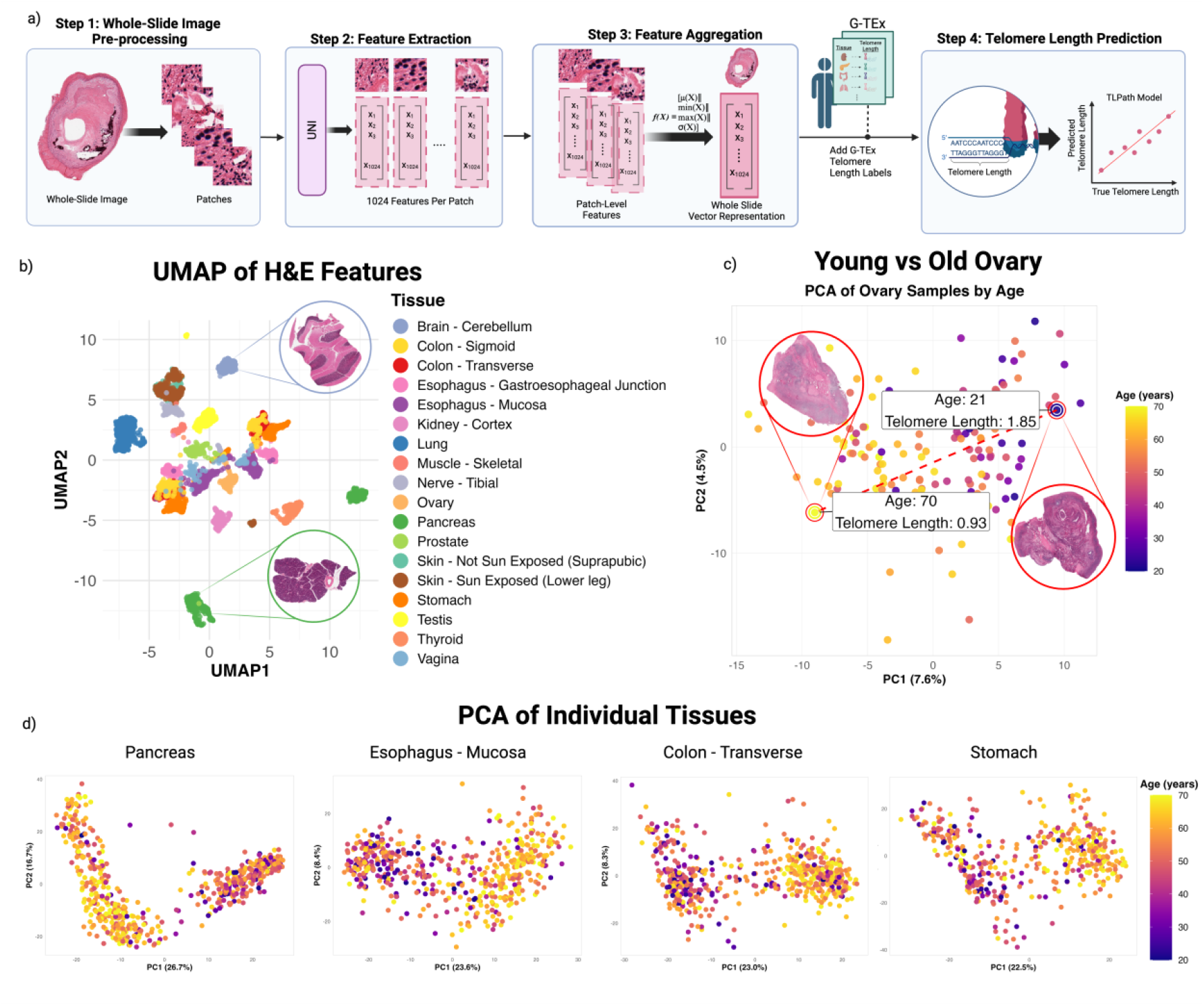
TLPath framework for bulk-telomere length prediction from H&E images. (**a**) The TLPath pipeline begins with whole-slide H&E images, which are processed into patches (Step 1). The UNI model extracts 1024-dimensional feature vectors per patch (Step 2). These are mean aggregated into a 1024-dimensional whole-slide vector representation (Step 3). Finally, this representation is used to build a supervised linear model to predict tissue telomere length (Step 4). (**b**) A UMAP of morphological features per slide extracted from H&E-stained tissue sections. Each point represents a single tissue sample, colored by tissue type. Whole-slide image of biopsies of pancreas and brain – cerebellum is shown. The clustering patterns reveal tissue-specific architectural features. (**c**) PCA Visualization of all H&E features of the ovary colored by age (purple – young, and yellow – old). Whole-slide images of the two biopsies are shown. (**d**) PCA visualization of four individual tissues (pancreas, esophagus-mucosa, colon – transverse and stomach), with points colored by patient age. The first two principal components reveal age-related patterns in morphological features, with varying degrees of age-associated structure across different tissue types. The four examples shown show that H&E features can capture age-related differences in young (purple gradient) vs old (yellow gradient).

**Figure 2:**
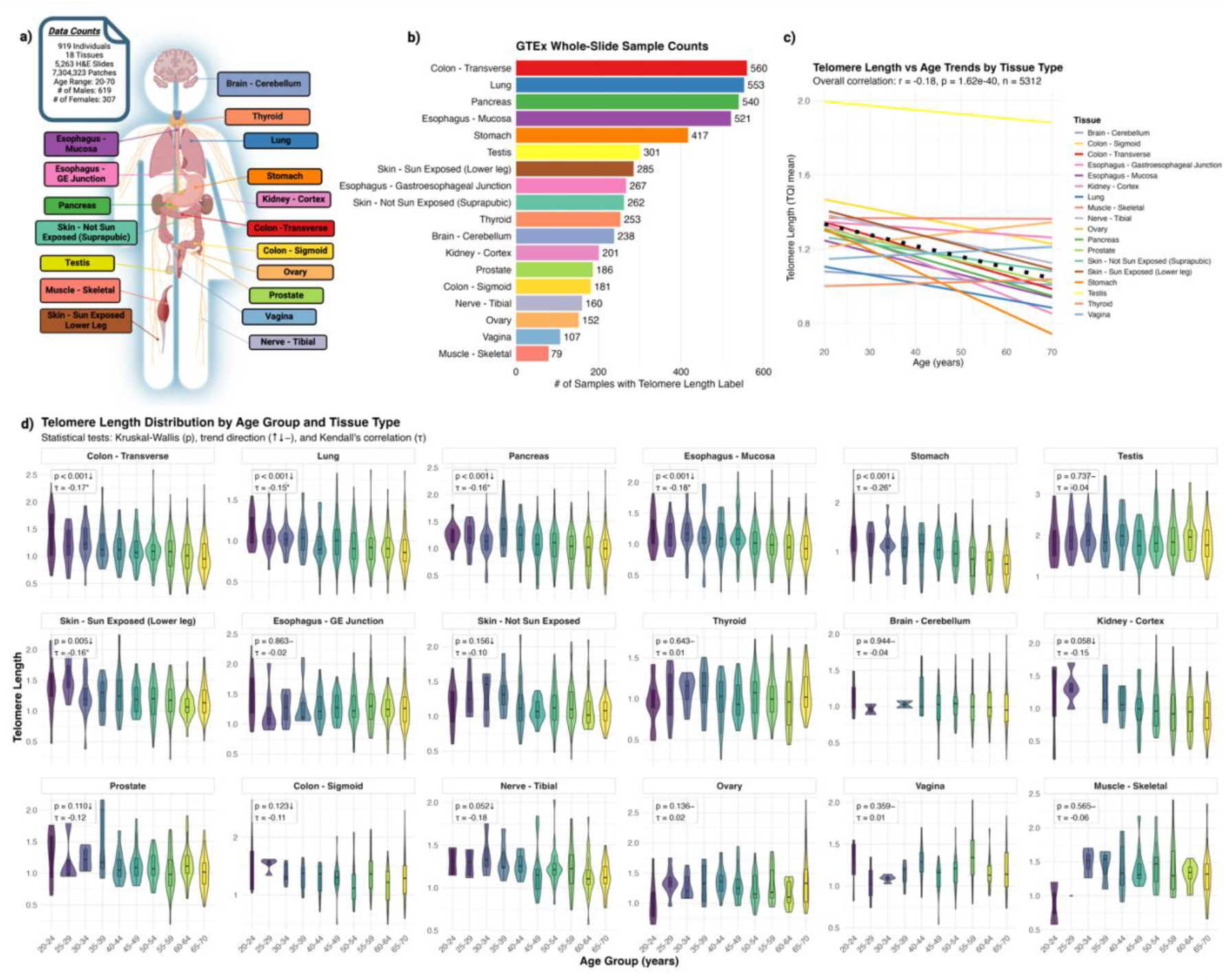
Overview of GTEx bulk-telomere data. **a)** The tissue types included in our study are from 919 individuals across 18 tissue types, totaling 5,263 H&E slides. **b)** Sample counts per tissue type in the GTEx dataset. **c)** Trendline of all 18 tissues of telomere length vs. age on a single plot. Testis shows the highest telomere length with slight decrease across age. Overall, there is a negative correlation with telomere length and age, with some tissues showing a positive correlation such as ovary and vagina. **d)** Visualizing relative telomere length levels in 18 different tissues, with subjects grouped into 5-year age intervals (20-64) and a final group of 65-70 years. Significant results (p < 0.001) include colon – transverse, lung, pancreas, esophagus – mucosa, and stomach which show decrease of telomere length as age groups get older.

TLPath pipeline starts by *(Step 1: <u>Preprocessing and Normalization</u>)* preprocessing the given whole-slide images (WSIs) by removing the background, color normalization, and segmentation into 512×512-pixel patches, where low-tissue content patches are removed, as described in **Methods 4.1**. There are 1,387 patches on average per whole-slide images across all 18 tissues. *(Step 2: <u>Feature Extraction</u>)* TLPath then extracts morphological features of each patch using a pre-trained pathology foundation model, UNI, a Vision Transformer (ViT) architecture trained on the more than 100K pathology WSI. This yields 1,024-dimensional feature vectors from each patch and is called – a patch-level embedding. (*<u>Step 3: Generating Whole-Slide Embedding</u>*) While our morphological features are available for each patch, there is only one telomere label per biopsy. This creates a gap between the resolution of telomere labels (one per slide) and features (patch-level). We therefore create a whole slide-level morphological embedding by aggregating across all patches of a slide using statistical measures, mean, yielding a 1,024-dimensional slide-level feature vector, detailed in **Methods 4.2**. Demonstrating the biological relevance of this representation, we observed that this slide-level representation can: 1) capture the tissue-type specific morphologies and can differentiate different tissue types (**Figure 1b**) and their prior known relationship (**Figure S1**), and 2) capture tissue deterioration in tissue during aging, demonstrated by separating tissues of different age with no prior training, for most tissue (**Figure 1c**). A few examples are illustrated in **Figure 1d** including colon, pancreas, esophagus, and stomach, where all tissues are provided in **Figure S2**.

(*Step 4: <u>Telomere Length Prediction</u>*) In the final step, we used the slide-level morphological features to build a supervised model (random forest) to predict bulk-telomere length labels available from the GTEx project. The model is trained and optimized using nested cross-validation to ensure robustness, with hyperparameter tuning step described in detail in **Methods 4.3**. To provide a baseline comparison, chronological age is used as an alternative predictor. The final prediction performance is measured using a Pearson’s correlation (r), between true and predicted bulk-telomere length (**Methods 4.4**).

#### Training and Testing Strategy

TLPath is trained separately for each tissue type, considering differences in their morphology, for 18 tissues where paired H&E images and telomere length labels are available (n > 5,263, GTEx). The number of samples per tissue for tissue types used for TLPath is provided is **Figure 2b**, where the colon, lung, pancreas and esophagus are among tissues with most samples. In most tissues, telomere length decreased with age as trends shown in **Figure 2c**, suggest the decrease in TL is strongest in colon and no age-related change in the testis. We provided their cross-tissue relationship in **Figure S3**. This is consistent with original study analyzing the telomere data (10) and other reports (14, 15), where testis have longer telomeres due to high telomerase activity and being immune-privileged, whereas stomach goes through consistent damage throughout lifespan. Telomere length distribution in five-year groups for each tissue is provided which shows a general trend of decrease in bulk-telomere length with age. Although, the pattern isn’t universal - some tissues like thyroid (p = 0.64, τ = 0.01) and vagina (p = 0.36, τ = 0.01) show less consistent age-related decline, and testis (p = 0.74, τ = -0.04) shows relatively stable lengths across age groups **Figure 2d** for further visualization. TLPath is evaluated in the GTEx cohort for each tissue separately, using a nested cross-validation approach, with 5 outer folds that the model has never seen for performance estimation and 3 inner folds for hyperparameter tuning, to assess the model’s ability to predict telomere length from H&E images across GTEx tissues. TLPath’s features relationship with biological and technical covariates in computed and provided in **Figure S4-6**.

### 2.2. TLPath Enables Age-Independent Telomere Assessment

Before we assess TLPath performance, we establish three baselines for telomere length determination: telomere length correlation between technical replicates (r = 0.91, derived (10)), cross-assay correlation between Southern blot vs. Luminex (r = 0.7, derived from (10)), and chronological age predictive power, which showed limited correlation (r = 0.21) across all 18 tissue types, peaking in stomach (r = 0.39, **Figure 2d, 3a**). TLPath’s morphological features demonstrated substantially stronger predictive capability than age, achieving an average correlation of 0.32 across tissues, with >0.4 correlation in 11 out of 18 tissue types. Notably, tissues with >300 samples showed markedly higher performance (r = 0.51) compared to those with fewer samples (mean r = 0.22), exemplified by pancreas (540 samples, r = 0.66) and stomach (417 samples, r = 0.57) (**Figure 3b**). Ovary (r = -0.12), vagina (r = -0.14), and colon-sigmoid (r = -0.05) exhibited negative correlation coefficients, likely due to insufficient sample sizes, as prior studies (16, 17) have consistently demonstrated that telomere length shortens with increasing age. TLPath trained using subset of samples without post-mortem artifacts including autolysis and with no pathologist-pointed pathologies (N=2,604), further improved our model’s performance in above well-represented tissues (**Figure S8**).

**Figure 3:**
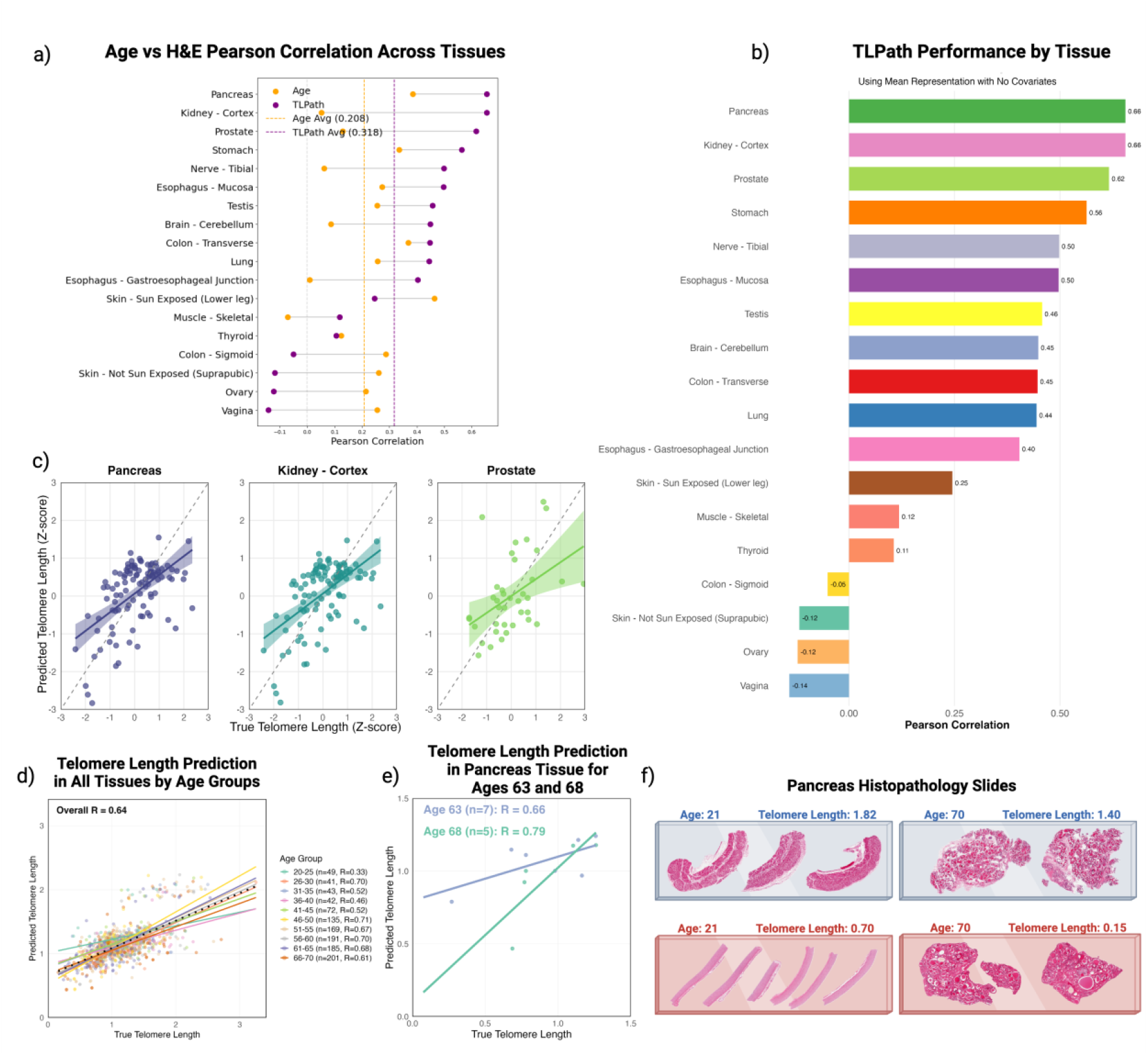
TLPath performance assessment and comparison against chronological age. **(a)** Comparison of Pearson correlations between chronological age (Linear Regression model) and H&E features (TLPath with H&E features as input), shown across different tissue types. Dashed lines indicate average correlations for age model (mean r = 0.21) and TLPath (mean r = 0.32). **(b)** TLPath’s performance in predicting telomere length across tissue with significance: there are 11 tissues in which TLPath can predict the telomere lengths with correlations >0.4. Additionally, there are 4 tissues (colon – sigmoid, skin – not sun exposed, ovary, and vagina) where TLPath has no predictive ability (correlation r < 0). **(c)** The strongest correlations were observed in pancreas (r = 0.66, n = 540), kidney – cortex (r = 0.66, n = 201), and prostate (r = 0.62, n = 186) and the scatter plots show each predicted vs. true telomere length (Z-scored) values for these tissues. The darker regression lines indicate the best fit line. **(d)** Telomere length prediction by age group in all tissues, with number of samples shown. The overall correlation (black-dotted line) shows a positive correlation between true vs predicted telomere lengths. **(e)** Telomere length prediction in pancreas tissue in age-matched samples of age 63 with 7 individuals and age 68 with 5 individuals. Both show positive correlation in predicting true vs predicted telomere length with average correlation 0.66 and 0.79 in age 63 and 68 respectively. **(f)** Pancreatic histopathology comparing samples from a young (Age 21, left slides) and old (Age 70, right slides) individual, representing extremes of age and telomere length within the study cohort. Blue slides: Samples with longer telomere lengths. Red slides: Samples with shorter telomere lengths. This figure visually contrasts tissue morphology, highlighting potential age-related changes and the association between shorter telomeres and altered pancreatic tissue structure.

Critically, TLPath’s real-world utility was validated through rigorous age-matched analyses. First, the model maintained significant predictive power when comparing samples from individuals within the same 5-year age bracket, demonstrating ability to distinguish between short and long telomeres independently of age-related variation (**Figure 3d, S7**). This capability was further validated at higher resolution, where TLPath successfully identified telomere length differences between individuals of the exact same chronological age (**Figure 3e**). Notably, the model accurately identified cases that deviate from typical age-telomere length relationships - detecting both young individuals with unexpectedly short telomeres and elderly individuals maintaining longer telomere lengths (**Figure 3f**). This age-independent predictive capability was consistently observed across seven diverse tissue types, suggesting TLPath captures fundamental morphological signatures of telomere length beyond age-associated changes. These results demonstrate TLPath’s potential for identifying telomere length variations within age-matched cohorts, a critical capability for applications.

### 2.3. TLPath uses senescence-like morphology to determine telomere length

To interpret TLPath, we first identified the top five most important features for the decision-making process focusing on pancreas (best performing tissue) based on model’s SHAP’s value (**Figure 4a, S8**). To link these deep-learning abstract features to interpretable histological patterns, we identified high-activation patches of each feature from slides with highest and lowest telomere length (**Figure 4b**, 15 slides from each group). This enabled systematic visualization of the tissue morphological patterns captured by these features. We interpreted these features in two parts: identifying simple cellular features and then higher-order pathology features (**Figure 5a**). For the first part, we quantified cell and nucleus-level features in above patches using QuPath (18), a widely used quantitative pathology tool, yielding 14 properties (**Table S1**). This revealed that mean nucleus-to-cell ratio, a morphological marker of senescence (19), is higher in patches from shorter telomere samples (**Figure 5a-c**). This is consistent across multiple features tested (**Figure S9**). Conversely, variation in nuclear intensity and nuclear circularity were higher in patches with longer telomeres. This suggests that TLPath might be capturing indirect short telomere induced cell senescence to determine the telomere length. We next identified higher-order features of these patches using a vision-language model pre-trained with large number of paired slide and pathology reports. This unique training provides it an ability to determine a probability to pathology terms that are the closest associated with a given patch. We applied the recently developed method CONCH on above patches (detailed steps in **Methods 4.6.2**) revealing that morphological features: necrosis and fascia were higher in samples with short telomere length, whereas hypertrophy is the only term higher in long telomere length samples. (**Figure 5c and S10-11**).

**Figure 4:**
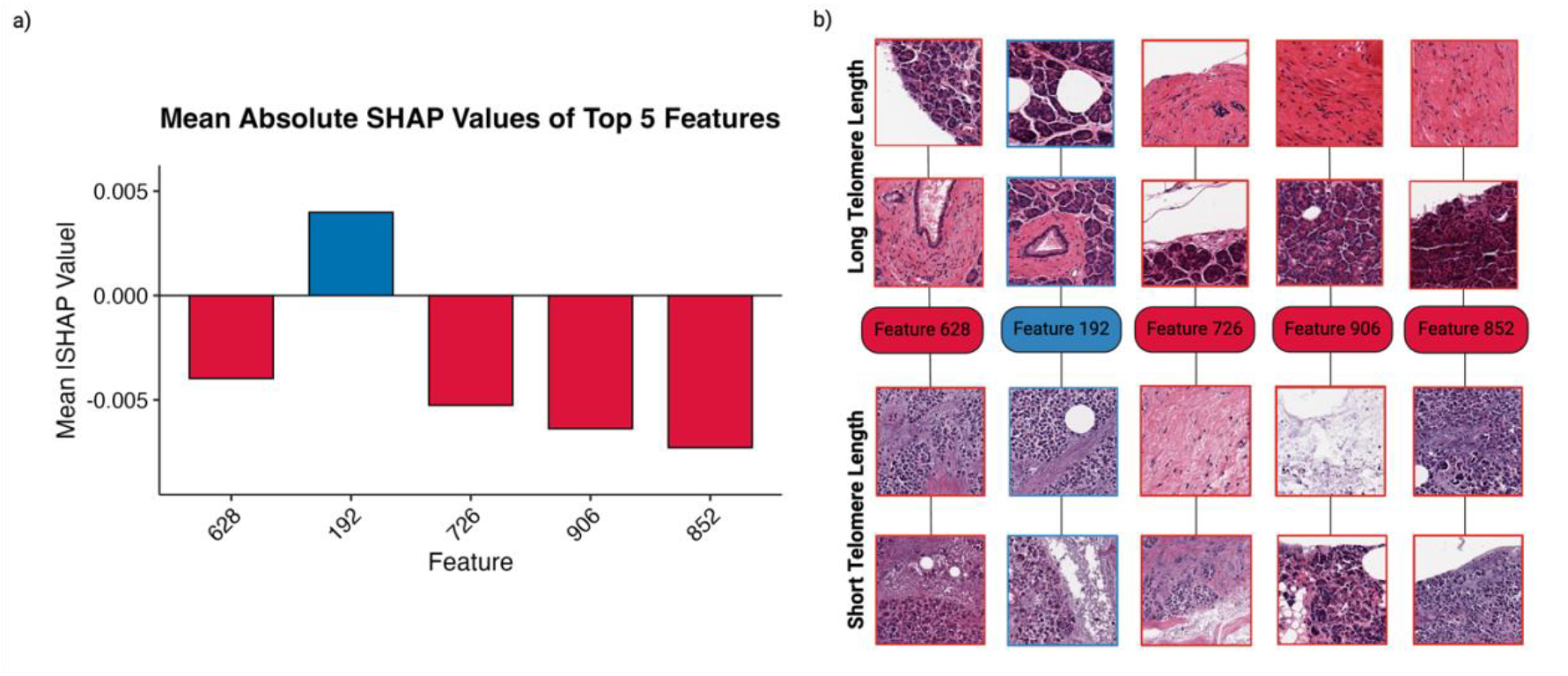
Top 5 features for TLPath and corresponding short and long telomere length patches. (**a**) The SHAP (SHapley Additive exPlanations) value analysis identified UNI feature 628, 192, 726, 906 and 852 as top five TLPath features with direction of contribution, to its decision making, focused on pancreas. Blue indicates features positively correlated with telomere length, and vice-versa. (**b**) Illustrative example patches for the top five features for Pancreas. The patches located on the top have longer telomere lengths, and the bottom patches have shorter telomere lengths.

**Figure 5:**
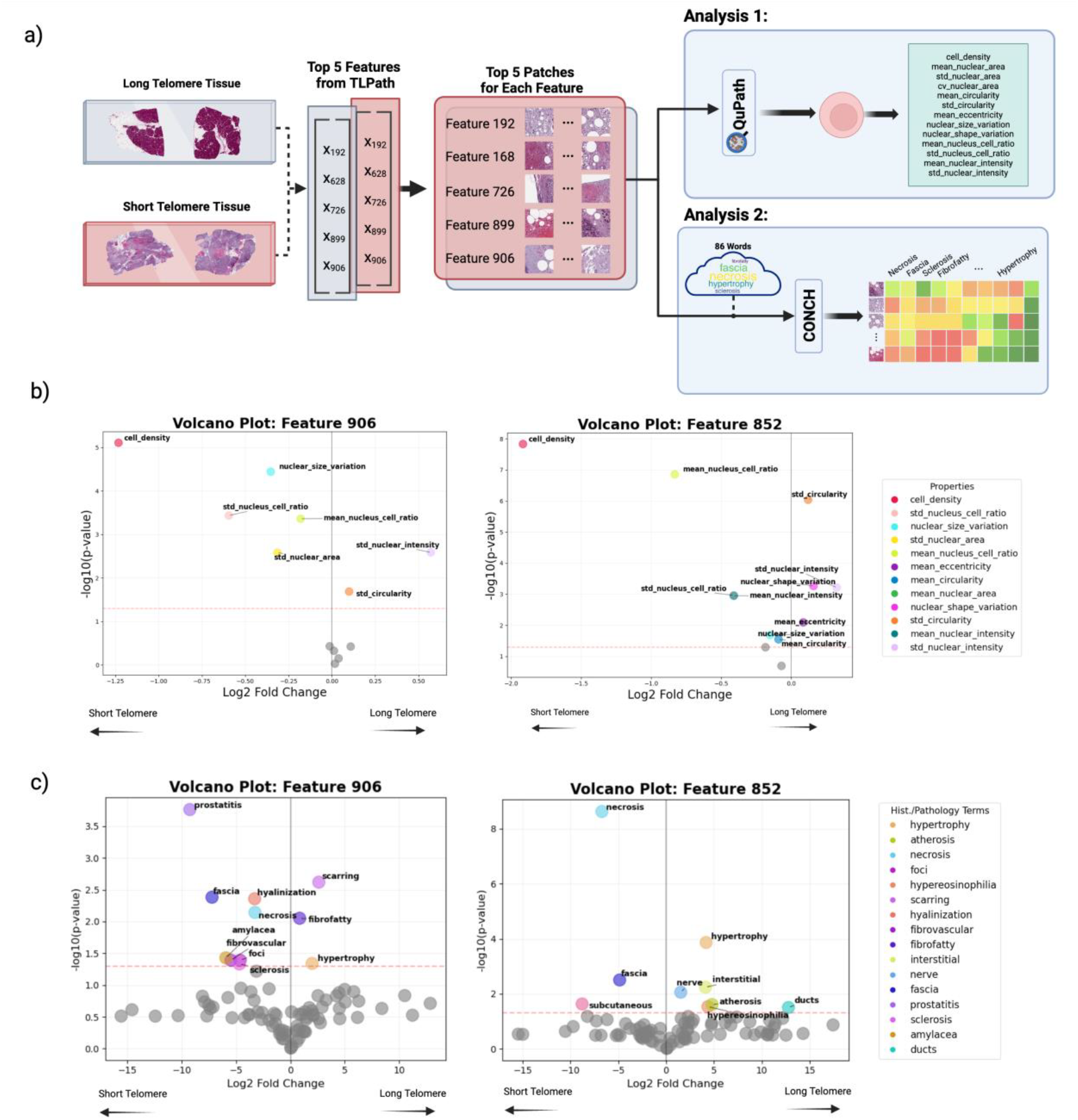
Mechanistic interpretation of TLPath features. **(a)** High-activation patches of TLPath’s most important features are extracted from long and short telomere samples. This was fed into QuPath (analysis 1) to identify cells and measure 14 distinct cellular and nuclear morphological properties (**Table S1**) and CONCH model (analysis 2). The average confidence score is calculated for all terms, across all patches. **(b)** The plot compares the long versus short telomere patches for features 906 and 852, depicting log fold changes (x-axis) against statistical significance (−log10 p-value, y-axis). Points right of zero indicate properties elevated in long telomere samples, while points left of zero show properties enriched in short telomere samples. The plots highlight differential cellular morphology characteristics between long and short telomere conditions. (**c**) The plots highlight the association of our terms from GTEx pathology reports to features patches 906 and 825, with long and short telomere from WSI.

### 2.4. Comprehensive benchmarking of TLPath steps to design an optimal architecture

To develop a robust framework for telomere length prediction from H&E images, we systematically evaluated each component of TLPath’s architecture through comprehensive benchmarking across diverse tissue types. Our analysis spanned different feature extractors and aggregation strategies, as well as modeling approaches to identify the optimal combination that maximizes both prediction accuracy and cross-tissue generalizability. TLPath integrates three key components that were systematically benchmarked for performance across all tissue types: 1) patch-to-slide aggregation through various pooling strategies (TITAN’s attention-based for CONCH features and mean, min-max, concatenated features for UNI), 2) Supervised model selection (random forest, linear regression, support vector regression, elastic net, ridge regression, and lasso regression) for telomere prediction and 3) Co-variate inclusion (age, sex, BMI, ethnicity).

Firstly, the mean feature aggregation method overall out-performs other whole-slide representation methods. Secondly, when examining strictly common tissues where all approaches could generate predictions (**Figure 6a**), elastic net achieved marginally higher mean correlation (r ≈ 0.50) compared to second best random forest (r ≈ 0.49). However, this comparison masks a critical difference in model versatility: while elastic net could only generate predictions for 10-12 tissues depending on the aggregation method, random forest successfully generated predictions across all 18 tissue types (**Table S2**). Performance of different models for the top three tissues are provided in **Figure 6b**. Finally, we note that adding biological covariates of age, sex, BMI, and ethnicity, only minorly improved TLPath model, suggesting that H&E morphology features might already be capturing these covariates already.

**Figure 6:**
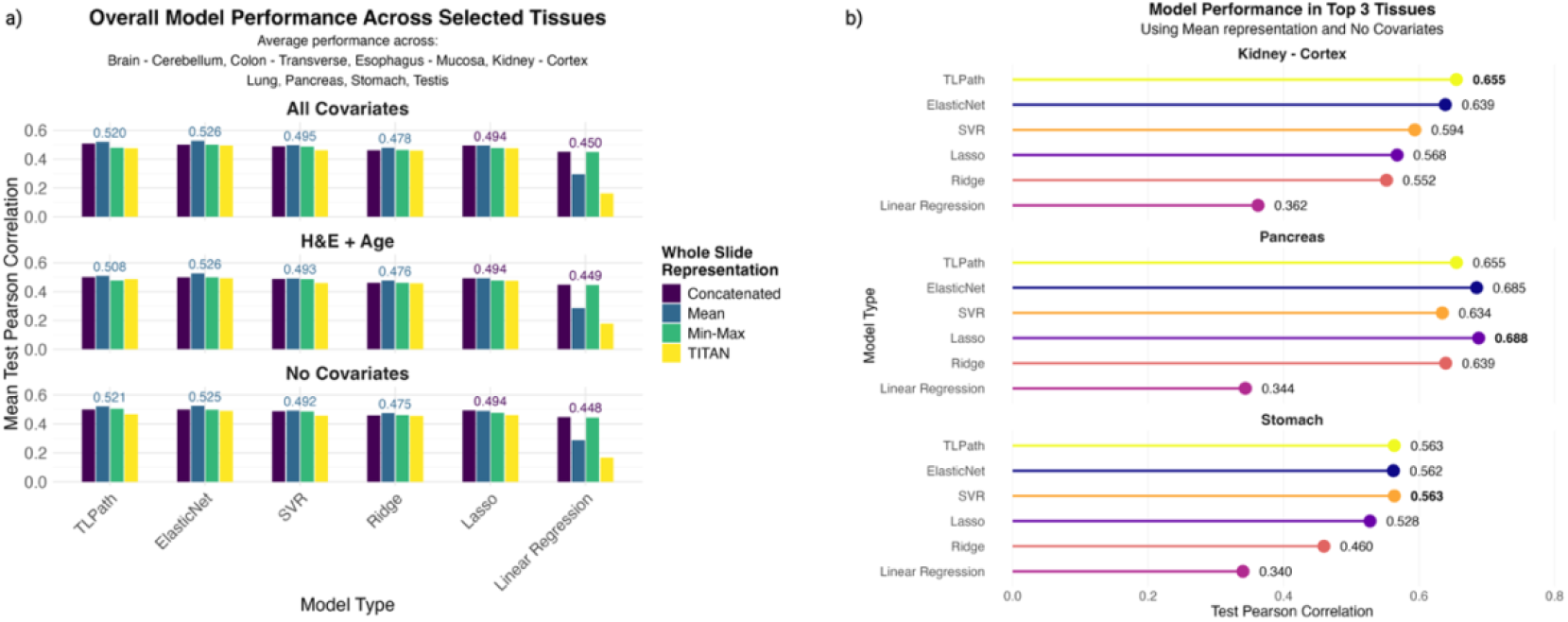
Comprehensive model performance benchmarking. (**a**) Performance comparison across six model types (TLPath, Elastic net, SVR, Ridge, Lasso, and Linear regression) using four different whole slide representation strategies (mean, min-max, concatenated, and TITAN). Results are shown for three different covariate configurations: All Covariates (top), No Covariates (middle), and Age Only (bottom). Performance is measured using Pearson correlation coefficients averaged across strictly common tissues where all models could generate predictions. (**b**) Detailed model performance comparison for the top 3 performing tissues (pancreas, kidney - cortex, and stomach) using the mean representation with no covariates. For each issue, performance is shown across different model types (TLPath, elastic net, linear regression, ridge, lasso, and SVR), Notably, while TLPath and elastic net show comparable performance in these tissues, TLPath maintains predictive capability across all tissue types, whereas other methods are limited to a subset of tissues.

### 2.5. Predicted bulk-telomere length is associated with T1/2 Diabetes across multiple tissues

Using medical history data from GTEx donors, we next examined the association between predicted telomere length and various diseases across different tissues while controlling for confounding factors such as age and sex. We focused on samples lacking measured telomere (N = 2,800), where the disease-specific cases and the available number of tissue biopsies for testing are provided in **Figure 7A**. We first observed an overall trend, although weak, of individuals with chronic diseases with shorter telomere length vs. ones who do not. However, this association is strongest for Type 1 and Type 2 diabetes (present in 62 and 217 donors, respectively) demonstrated consistent negative associations – shorter telomere in individuals’ disease across all tissues. We note colon– transverse tissue examples with statistically significantly lower telomere length (both true and predicted) in individuals with T2D vs. not (β = -1.46, P < 0.05, **Figure 7A-B**). These associations are then examined on the measured telomere (N= 8079). We observe the associations are consistent and coefficients are magnified in *measured* telomere length, except for kidney – cortex (β = -8.28, P < 0.05) with type 1 diabetes (present in 54 donors). We note that lungs are affected by non-respiratory diseases such as hypertension and rheumatoid arthritis, however further investigation is warranted. (**Figure 7A-B, *left panel***).

**Figure 7:**
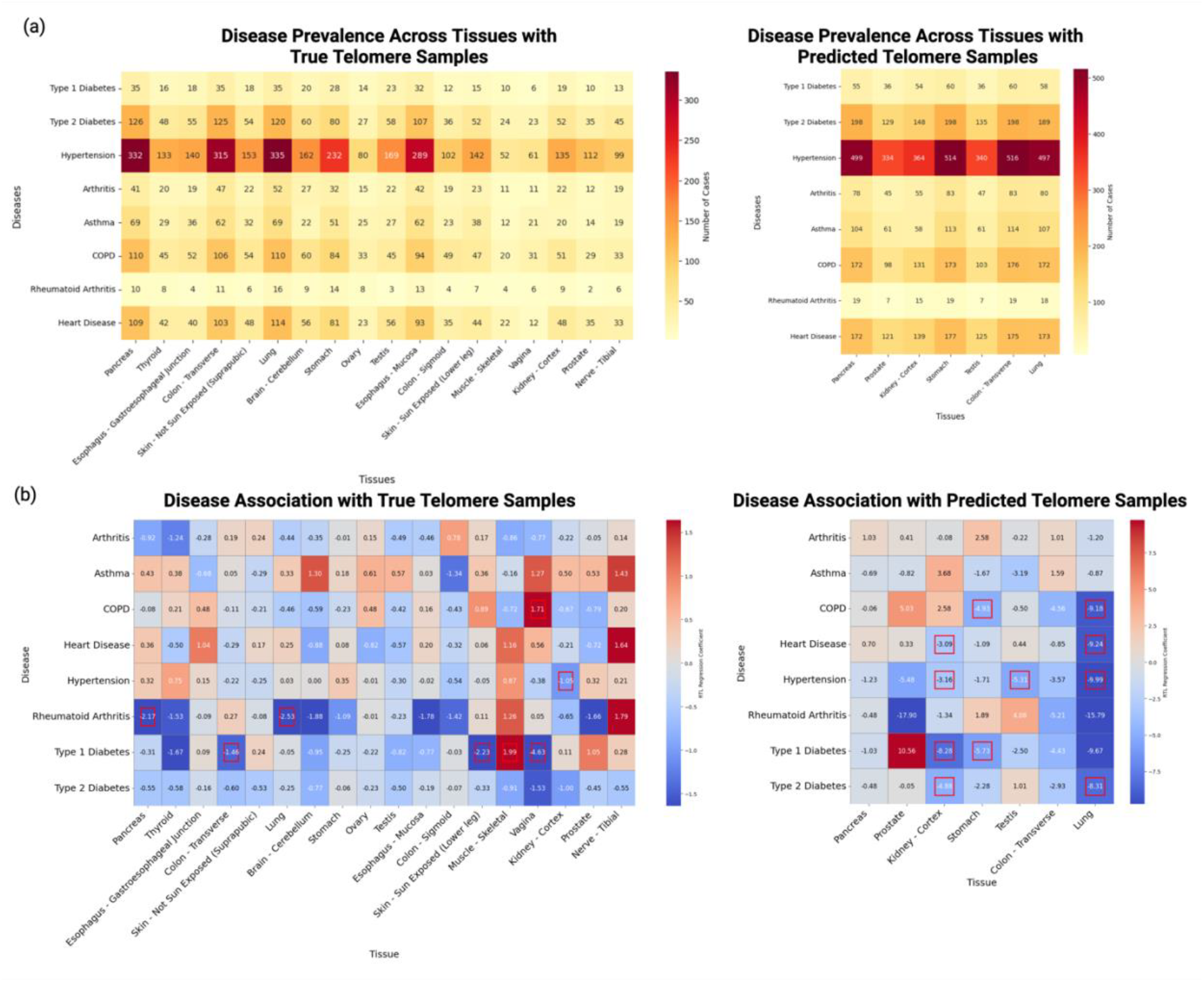
Disease associations with telomere length across GTEx tissues. (**a**) Disease prevalence heat maps showing the distribution of cases across tissues in both true telomere samples (left, 18 tissues) and predicted telomere samples (right, 7 tissues). Hypertension shows the highest prevalence across both datasets (61-335 cases in true samples; 324-697 cases in predicted samples), followed by Type 2 diabetes (23-126 cases in true samples; 129-198 cases in predicted samples). Other diseases such as rheumatoid arthritis show lower prevalence across all tissues. (**b**) Heat maps of regression coefficients demonstrating the association between diseases and telomere length while controlling for age and sex. The true telomere samples (left) show more modest associations with coefficients generally ranging from -4 to 4, while predicted telomere samples (right) display stronger associations ranging from -20 to 20. Type 1 and Type 2 diabetes consistently show negative associations across tissues in both datasets, with significant associations (marked with asterisks, p < 0.05) observed particularly in colon-transverse tissue for Type 1 diabetes (β = -1.46) in true samples.

## 3. Discussion

We here present TLPath, a digital pathology framework trained on >7 million images to predict normal tissue bulk-telomere length across more than ten human tissues from routinely available H&E-stained histopathology images. Our method demonstrates the potential of capturing aging processes directly from high-resolution H&E-based tissue images. To our knowledge, this is the first approach capable of predicting relative telomere length from H&E slides, offering a new window into telomere research through machine learning analysis of tissue morphology. TLPath outperformed chronological age in predicting telomere length, demonstrating H&E-based morphology’s ability to capture individual specific telomere information. Mechanistic interpretation of the model revealed that TLPath determines short-telomere samples by using senescence-like cell morphology (high nuclear-to-cytoplasmic ratio) along with tissue damage (necrosis) and fascia-like structures. TLPath’s performs better in tissue types with more training data, suggests that with more samples, TLPath can become more accurate in additional tissues.

TLPath faces three key limitations. First, a fundamental limitation relates to the specificity of TLPath for telomere length versus other aging hallmarks. Many hallmarks of aging are correlated and produce similar morphological changes at the tissue and cellular level. The senescence-like morphology and tissue damage patterns that TLPath identifies may reflect not only telomere shortening but also related processes like epigenetic changes, chronic inflammation, and protein misfolding. Unlike direct measurement methods, TLPath indirectly predicts telomere length through visible manifestations of cellular aging. While this approach may capture functionally relevant changes that impact disease, it cannot definitively determine whether the morphological features detected are directly caused by telomere shortening or represent broader senescence signatures that correlate with shortened telomeres. Additionally, it fails to detect telomere shortening that has not yet produced visible tissue alterations.

Second, TLPath likely cannot distinguish between shortened telomeres and telomere dysfunction independent of length. Given the accumulating importance of telomeric damage/dysfunction (2), this represents a significant limitation in fully characterizing telomere biology through morphological features alone. Furthermore, TLPath cannot distinguish between different causes of telomere shortening, such as replicative exhaustion versus accelerated shortening due to environmental stressors or genetic factors. Third, TLPath provides only bulk-tissue level predictions, averaging telomere lengths across the entire slide and masking both cell-type specific and spatial variations. This limitation prevents assessment of the critical shortest telomeres within individual cells (2, 8). These limitations collectively highlight that while TLPath offers valuable insights into telomere biology at scale, it should be interpreted as part of a broader toolkit rather than a replacement for specialized molecular analyses when mechanistic specificity is required.

Further, future studies should focus on developing tissue-specific models incorporating information at cell level resolution and labels at cell-type resolution and extending this model to mice studies where large-scale archives are available and interventions are tested with tissue biopsies more readily available. However, extension to mice is challenging considering the longer telomere length in mice vs. human and its shortening dynamics. Finally, while this study shows the utility of computational pathology for predicting aging biology, focused on telomere length here and its extension to other aging process will be of high reward, including senescence, chronic inflammation, genomic instability.

## 4. METHODS

### 4.1. sData Acquisition

Whole-slide images were obtained from the GTEx public portal via v10 release (https://www.gtexportal.org/home/). All slides were digitized at 20X magnification (∼0.5 microns per pixel) using an Aperio scanner (SVS format). Protected data, including participant demographic information (chronological age), lifestyle factors, clinical information, and gene expression data, were accessed through dbGAP (Request accession ID: 39103) (20). Telomere length measurements (TQImean) were created in the Institute for Population and Precision Health (IPPH) Laboratory at the University of Chicago were obtained from the publicly available GTEx dataset (https://www.gtexportal.org/home/downloads/egtex/telomeres).

Telomere length measurements were obtained from GTEx project using a Luminex-based assay that measures bulk-tissue relative telomere length (RTL) and described at (28). Briefly, RTL is defined as the telomere repeat abundance in a sample relative to a standard reference DNA sample. From an initial set of 7,234 tissue samples from 962 donors, 6,391 tissue-specific RTL measurements from 952 donors were retained after quality control, which included removal of failed measurements and within-tissue outliers. The Luminex assay was validated with orthogonal assays. This provided ground truth relative telomere length measurements across 18 different tissue types, enabling development of TLPath.

### 4.2. Dataset Selection and Filtering

While the GTEx project encompasses samples from 54 different tissues collected from nearly 1,000 adult post-mortem donors, we focused our analysis on a subset of tissues based on two key criteria:

1. Availability of H&E-stained tissue images
2. Number of samples >70 for robust training

The threshold of at least 70 samples provides model enough data to train and fine-tune on, given our nested cross-validation training. These selection criteria led to the exclusion of 36 tissue types, such as whole blood (lacking H&E slides) and tissues with insufficient sample sizes. The final dataset comprised 5,263 samples across 18 distinct tissues.

### 4.3. Whole-slide image processing and feature extraction

We implement and compare the vision encoders UNI (21) and TITAN (22) for WSI feature extraction. While TITAN gives us whole slide representation, UNI gives patch level embeddings that we then aggregate to create whole slide representations.

UNI is a large vision transformer (ViT), pretrained using the state-of-the-art self-supervised learning algorithm DINOv2(23) - a student-teacher knowledge distillation for ViT architecture. UNI was pretrained on the Mass-100K dataset, a dataset containing more than 100 million tissue patches across 20 major tissue types. To preprocess whole slide images for UNI, we implemented Sobel edge detection (24) to first identify regions of the slide containing tissue. Each 20x magnification WSI was then segmented into 512 × 512-pixel patches. Patches were excluded if more than 50% of their pixels exhibited low weighted gradient magnitude, as determined by Sobel. RGB color normalization was applied to remaining tiles using Macenko’s (25). Following image preprocessing, these patches are passed to UNI to extract 1024-dimention patch level features. We create a whole slide representation by aggregating each 1,024-patch level feature together. There are 3 different ways in which we aggregate the patch level features from UNI to get slide level representation mentioned (See **5.41**).

We compare our WSI aggregated features from UNI against TITAN, a state-of-the-art WSI vision encoder. TITAN was pretrained on 335K WSI across 20 major tissue types. To process whole slide representation from TITAN, whole slide images were preprocessed using a multi-step pipeline available in CLAM (26). First, tissue regions were segmented from the background using color thresholding in the HSV color space, followed by median filtering and binary thresholding. The segmented tissue regions were then filtered to remove small artifacts and holes using contour area-based filtering. From the identified tissue regions, non-overlapping patches of size 448×448 pixels were extracted at the highest magnification level (patch level = 0). The extraction process used a padding mechanism to ensure complete coverage of tissue regions and employed a four-point contour checking method to verify if patches contained sufficient tissue content. These patches were then passed to CONCH_v1.5 (27) to extract 768-dimensional feature for each patch. The extracted features from each patch were arranged in a two-dimensional grid to preserve positional information and then given as input to TITAN. Finally, TITAN generated a 768-dimentional WSI feature representation.

### 4.4. TLPath Training

#### 4.4.1. Data Preparation

To ensure consistent feature scales, our input features, including the aggregated UNI features from Whole Slide images and clinical covariates (age, sex, race or ethnicity, BMI and smoking status), were normalized using StandardScaler from scikit-learn. The scaler was fitted exclusively on the training data to prevent data leakage, and the resulting transformation was subsequently applied to the validation and test datasets.

#### 4.4.2. Model Development

To predict telomere length from the extracted H&E Whole Slide Image features and matching clinical covariates, we implemented and evaluated a suite of supervised models with regularization approaches. For a baseline performance assessment, we developed two reference models. First, a standard multiple linear regression model was implemented to establish a direct mapping between our input features and telomere length without regularization. Second, we constructed a univariate linear regression model using only chronological age as a predictor to serve as a clinical baseline, enabling comparison with current practice standards.

To address the challenges of potential overfitting and feature collinearity, we implemented lasso regression (L1 regularization) and ridge regression (L2 regularization). The first approach was selected to help identify the most relevant input features by shrinking less prominent features’ coefficients to zero and second to handle multicollinearity among features by imposing a penalty on the magnitude of coefficients. As a combination of the two, we also implemented elastic Net (L1 + L2 regularization).

Additionally, we implemented and evaluated two non-linear supervised regression approaches: support vector regression (SVR) and random forest regression. The SVR model was implemented with both linear and RBF kernels to capture potential non-linear relationships in the data. The Random Forest model was employed as an ensemble learning method to capture complex patterns while providing feature importance measures. Hyperparameters for all models were optimized using k-fold cross-validation to ensure robust performance estimation (See **4.4.3**).

#### 4.4.3. Training Strategy

We employed a nested cross-validation framework to ensure robust model evaluation and optimal hyperparameter selection. The outer loop consisted of 5-fold cross-validation for model validation, while the inner loop employed 3-fold cross-validation for hyperparameter tuning. For the regularized models, we performed grid search over a comprehensive range of hyperparameters. The Lasso model’s α parameter was tuned over {0.001, 0.01, 0.1, 1, 10, 100}. The Ridge model’s α parameter was explored over {0.001, 0.01, 0.1, 1, 10, 100, 1000, 10000, 100000}.

For SVR, we performed grid search over the C parameter {0.1, 1, 10}, epsilon values {0.1, 0.2, 0.3}, and kernel types {linear, RBF}. For Random Forest, we employed randomized search with 20 iterations over various hyperparameters including number of estimators (50-200), maximum depth, minimum samples split (2-10), minimum samples leaf (1-4), and maximum features selection methods (‘sqrt’, ‘log2’).

To ensure the robustness of our results, we repeated the entire nested cross-validation process using multiple random seeds. This approach helped account for potential variability introduced by random data partitioning.

#### 4.4.4. Model Evaluation

We evaluated model performance using multiple complementary metrics. The primary metrics included Mean Squared Error (MSE) for assessing prediction accuracy, R-squared (R^2^) for explaining variance in telomere length, and Pearson correlation coefficient (r) for measuring the strength of the relationship between predicted and actual values. For the Pearson correlation coefficient, we calculated 95% confidence intervals using Fisher’s z-transformation to assess statistical significance.

The final model for each regression type was trained on the complete training dataset using hyperparameters optimized through nested cross-validation. These models were then evaluated on our test set, which remained unused during the training and optimization process, to obtain unbiased estimates of generalization performance.

### 4.5. Benchmarking

#### 4.5.1. Feature Aggregation Strategies

We systematically evaluated different feature aggregation strategies to get whole slide representation from features extracted via UNI to determine their impact on model performance:

- Mean Aggregation: Computing the average of feature vectors across all tiles
- Maximum and Minimum Aggregation: Selecting the maximum and minimum value for each feature dimension
- Combined Aggregation: Concatenating mean, minimum, max, and standard deviation Each aggregation strategy resulted in feature vectors of varying dimensions: mean aggregation resulted in 1,024-dimension feature vectors, maximum and minimum aggregation resulted in 2,048-dimension and the combined aggregation resulting in 4,096-dimension feature vectors.

#### 4.5.2. Evaluation Framework

We maintained the same nested cross-validation framework described in Section 4.3 for all benchmarking experiments. To ensure fair comparison, we evaluated each feature extraction and aggregation combination using all 6 regression models (Linear, Lasso, Ridge, Elastic Net, SVR and Random Forest). Performance evaluation was maintained the same as the model evaluation framework described in **Section 4.4** for all benchmarking experiments.

### 4.6. Interpretability of TLPath

To interpret TLPath’s decision-making process, we performed feature importance analysis using Random Forest weights on our 1024-dimensional feature vector. This analysis identified five key features (dimension #906, #852, #726, #192, and #628) that showed the highest SHAP values, indicating their strong influence on telomere length prediction (**Figure 5a, Figure S8**). To link these abstract features to interpretable histological patterns, we focused our analysis on pancreatic tissue, where TLPath showed the strongest performance. We selected 30 whole slide images representing extreme telomere lengths (top and bottom 15 telomere length slides). From each slide, we extracted the five patches showing the highest activation values (UNI-based) for each key feature, enabling systematic visualization of the tissue patterns captured by these features. Our subsequent analysis comprised two main components, detailed below.

#### 4.6.1. QuPath

To elucidate the morphological characteristics captured by TLPath, we leveraged QuPath’s (18) cell detection algorithms to quantify nuclear and cellular morphological features in WSI patches. We systematically analyzed 14 cellular properties, including nuclear metrics (area, perimeter), shape parameters (circularity, eccentricity), and nucleus-to-cell ratios. For each property, we computed both mean values and standard deviations across 150 patches derived from 30 pancreatic WSIs, equally divided between specimens with short and long telomere lengths.

This provided which morphological features unique to short vs. long telomere specimen. Median comparison was done using Mann-Whitney U test. Log2 fold changes were calculated using median values, with significance threshold set at p < 0.05. Properties were color-coded for visualization, and separate plots were generated for each feature to enable systematic comparison across all morphological parameters.

#### 4.6.2. Conch Interpretability

We explored the ability of CONCH (27), a vision-language contrastive model, to systematically associate visual features from H&E image patches with histological and pathological terms (n = 86). To do this, we first extracted the top 200 common terms from pathology categories and notes in our GTEx dataset, filtering for histology and pathology-specific terms. We then further filtered our terms, removing non-descriptive terms like “tissue”, “cells”, and “chronic”. Finally, we remove redundant words by keeping the root words and removing variations such as keeping “atrophy” and removing “atrophic”. This left 86 terms commonly used by pathologists that we used for our analysis (**Table S3**). We analyzed the average similarity confidence scores of CONCH for all terms across all patches, grouped by UNI feature activation and telomere length.

## 5. DATA AVAILABILITY STATEMENT

The data used in this study were obtained from the Genotype-Tissue Expression (GTEx) Project (accession: phs000424.v10.p2, Project titled #39103). The H&E histological images are publicly available and were accessed through the GTEx Portal v10 release (https://gtexportal.org/home/). Telomere length data were also obtained GTEx Portal (https://www.ncbi.nlm.nih.gov/gap/) Biological and Technical covariates, e.g. age, were extracted after appropriate institutional approval and data access agreement (General Research Use: phs000424.v10.p2.c1). All processed data, including derived tables and analyses generated in this study, have been deposited in Zenodo (ZENODO_DOI_PLACEHOLDER).

## 6. CODE AVAILABILITY STATEMENT

TLPath is available on GitHub as a software package for academic use. The complete source code and documentation can be accessed at: https://github.com/Sinha-CompBio-Lab/TLPath

## 8. ACKNOWLEDGEMENTS

This research was supported by the Sanford Burnham Prebys NCI-designated Cancer Center Start-up Fund for Sinha Lab and LEAP Fellowship for K.A.

## 9. AUTHORS CONTRIBUTION

S.S. conceptualized the idea of predicting telomere length from H&E. A.Y., S.S., K.A., developed the framework for predicting telomere length from H&E slides. A.Y. and K.A. wrote scripts of TLPath model. A.Y., A.A. and K.A. co-led the model interpretability analysis. K.A, A.Y. and A.A. developed independent methods to interpret the TLPath and identify decision-making features. A.Y, K.A. and A.A. wrote the initial manuscript with S.S. providing critical revisions.

## REFERENCES

1. Blackburn, E. H. (1991). “Structure and function of telomeres.” Nature, 350(6319), 569–573.

2. Rossiello, F., Jurk, D., Passos, J. F., & d’Adda di Fagagna, F. (2022). Telomere dysfunction in ageing and age-related diseases. Nature cell biology, 24(2), 135–147.

3. Hewitt, G., Jurk, D., Marques, F. D., Correia-Melo, C., Hardy, T., Gackowska, A., … & Passos, J. F. (2012). Telomeres are favoured targets of a persistent DNA damage response in ageing and stress-induced senescence. Nature communications, 3(1), 708.

4. Martínez, P., & Blasco, M. A. (2017). Telomere-driven diseases and telomere-targeting therapies. The Journal of cell biology, 216(4), 875.

5. López-Otín C, Blasco MA, Partridge L, Serrano M, Kroemer G. Hallmarks of aging: An expanding universe. Cell. 2023 Jan 19;186(2):243–278. doi: 10.1016/j.cell.2022.11.001. Epub 2023 Jan 3. PMID: 36599349.

6. Kimura, Masayuki, et al. “Measurement of telomere length by the Southern blot analysis of terminal restriction fragment lengths.” Nature protocols 5.9 (2010): 1596–1607.

7. Cawthon, R. M. (2002). Telomere measurement by quantitative PCR. Nucleic acids research, 30(10), e47–e47.

8. Norris, K., Walne, A. J., Ponsford, M. J., Cleal, K., Grimstead, J. W., Ellison, A., … & Baird, D. M. (2021). High-throughput STELA provides a rapid test for the diagnosis of telomere biology disorders. Human genetics, 140, 945–955.

9. Vera, E., & Blasco, M. A. (2012). Beyond average: potential for measurement of short telomeres. Aging (Albany NY), 4(6), 379.

10. Demanelis, Kathryn, et al. “Determinants of telomere length across human tissues.” Science, vol. 369, no. 6509, 2020, article eaaz6876. 10.1126/science.aaz6876.

11. Schildhorn, C., Jacobi, C., Weissbrodt, A., Hermstedt, C., Westhoff, J. H., Hömme, M., … & Melk, A. (2015). Renal phenotype of young and old telomerase-deficient mice. Mechanisms of Ageing and Development, 150, 65–73.

12. McDonough, J.E., Martens, D.S., Tanabe, N. et al. A role for telomere length and chromosomal damage in idiopathic pulmonary fibrosis. Respir Res 19, 132 (2018). 10.1186/s12931-018-0838-4

13. Kim, B., Ryu, KJ., Lee, S. et al. Changes in telomere length and senescence markers during human ovarian tissue cryopreservation. Sci Rep 11, 2238 (2021). 10.1038/s41598-021-81973-3

14. Chakradhar, S. (2018). Puzzling over privilege: How the immune system protects—and fails—the testes. Nature Medicine, 24(1), 2–5. 10.1038/nm0118-2

15. The paradox of longer sperm telomeres in older men’s testes: a birth-cohort effect caused by transgenerational telomere erosion in the female germline.” Molecular Cytogenetics 9.1 (2016): 12.

16. Ozturk S. The close relationship between oocyte aging and telomere shortening, and possible interventions for telomere protection. Mech Ageing Dev. 2024 Apr;218:111913. doi: 10.1016/j.mad.2024.111913. Epub 2024 Feb 1. PMID: 38307343.

17. O’Sullivan J, Risques RA, Mandelson MT, Chen L, Brentnall TA, Bronner MP, Macmillan MP, Feng Z, Siebert JR, Potter JD, Rabinovitch PS. Telomere length in the colon declines with age: a relation to colorectal cancer? Cancer Epidemiol Biomarkers Prev. 2006 Mar;15(3):573–7. doi: 10.1158/1055-9965.EPI-05-0542. PMID: 16537718.

18. Bankhead, P., Loughrey, M.B., Fernández, J.A. et al. QuPath: Open source software for digital pathology image analysis. Sci Rep 7, 16878 (2017). 10.1038/s41598-017-17204-5asd

19. Heckenbach, I., Mkrtchyan, G.V., Ezra, M.B. et al. Nuclear morphology is a deep learning biomarker of cellular senescence. Nat Aging 2, 742–755 (2022). 10.1038/s43587-022-00263-3

20. Mailman MD, Feolo M, Jin Y, Kimura M, Tryka K, Bagoutdinov R, Hao L, Kiang A, Paschall J, Phan L, Popova N, Pretel S, Ziyabari L, Lee M, Shao Y, Wang ZY, Sirotkin K, Ward M, Kholodov M, Zbicz K, Beck J, Kimelman M, Shevelev S, Preuss D, Yaschenko E, Graeff A, Ostell J, Sherry ST. The NCBI dbGaP database of genotypes and phenotypes. Nat Genet. 2007 Oct;39(10):1181–6. doi: 10.1038/ng1007-1181. PMID: 17898773; PMCID: PMC2031016.

21. Chen, R.J., Ding, T., Lu, M.Y. et al. Towards a general-purpose foundation model for computational pathology. Nat Med 30, 850–862 (2024). 10.1038/s41591-024-02857-3

22. Ding, Tong, Sophia J. Wagner, Andrew H. Song, Richard J. Chen, Ming Y. Lu, Andrew Zhang, Anurag J. Vaidya, et al. 2024. “Multimodal Whole Slide Foundation Model for Pathology.” arXiv preprint 2411.19666. https://arxiv.org/abs/2411.19666.

23. Oquab, Maxime, et al. “DINOv2: Learning Robust Visual Features without Supervision.” arXiv preprint 2304.07193 (2023). https://arxiv.org/abs/2304.07193.

24. Kanopoulos, N., Vasanthavada, N., & Baker, R. L. (1988). Design of an image edge detection filter using the Sobel operator. IEEE Journal of solid-state circuits, 23(2), 358–367.

25. Macenko, M., et al. 2009. “A Method for Normalizing Histology Slides for Quantitative Analysis.” Proceedings - 2009 IEEE International Symposium on Biomedical Imaging: From Nano to Macro, ISBI 2009, 1107–1110. 10.1109/ISBI.2009.5193250.

26. Lu MY, Williamson DFK, Chen TY, Chen RJ, Barbieri M, Mahmood F. Data-efficient and weakly supervised computational pathology on whole-slide images. Nat Biomed Eng. 2021 Jun;5(6):555–570. doi: 10.1038/s41551-020-00682-w. Epub 2021 Mar 1. PMID: 33649564; PMCID: PMC8711640.

27. Lu, M.Y., Chen, B., Williamson, D.F.K. et al. A visual-language foundation model for computational pathology. Nat Med 30, 863–874 (2024). 10.1038/s41591-024-02856-4

