## Supplementary Figures (S1-S13) + Supplementary Table 1 for "TLPath predicts telomere length in human tissues from histopathology images"

### Supplementary Figures and Tables

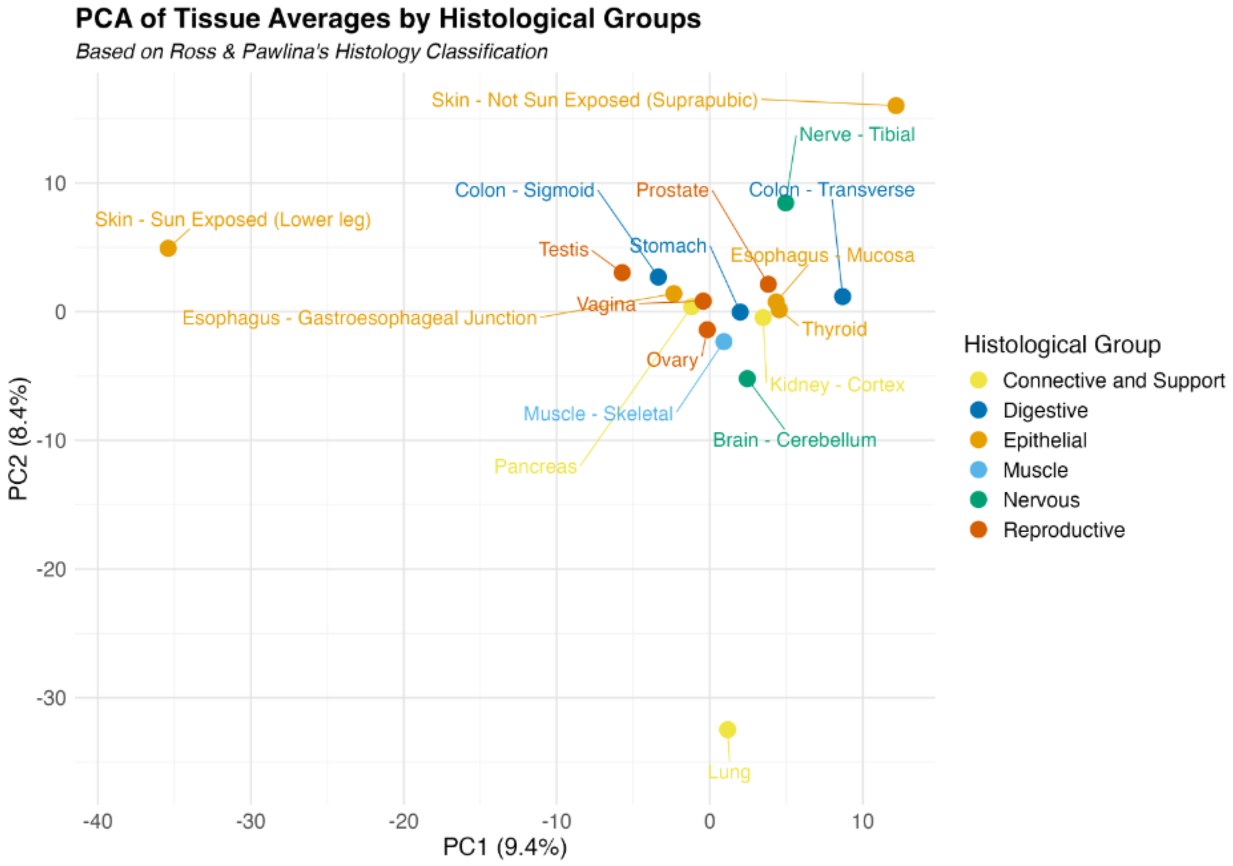


**Figure S1: Tissue relationships in principal component space**. Tissues are plotted according to their first two principal components derived from H&E image features. Tissues are color-coded according to Ross & Pawlina's tissue classification system: yellow for Connective and Support, blue for Digestive, orange for Epithelial, light nlue for Muscle, green for Nervous and red for Reproductive. The spatial arrangement reveals clusters of functionally related tissues such as testis, vagina, ovary and prostate with closely positioned points indicating tissues that share similar histological features. This visualization combines dimensionality reduction with similarity metrics to highlight both global tissue relationships and specific tissue-tissue connections.

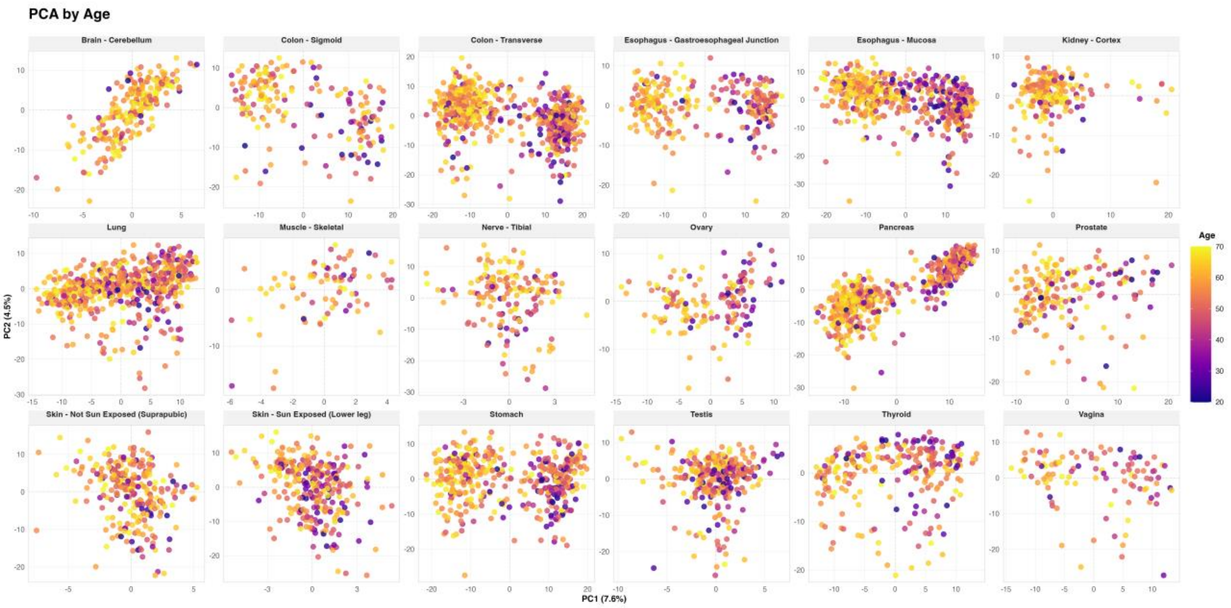


**Figure S2: Principal Component Analysis (PCA) of H&E image features reveals tissue-specific age-related patterns.** Scatter plots showing the first two principal components (PC1 vs PC2) of UNI-derived image features for 18 different tissues from the GTEx dataset. Each point represents a single sample, and colors indicate the donor's age (scale from purple [younger] to yellow [older]). The distribution and clustering patterns vary across tissues, suggesting tissue-specific relationships between morphological features and aging. Some tissues (e.g., colon - transverse, pancreas) show clear age-related gradients in the PCA space, while others (e.g., kidney - cortex, brain - cerebellum) display more diffuse patterns, highlighting the heterogeneous nature of age-related morphological changes across different tissue types. The variation in these patterns supports the need for tissue-specific approaches when analyzing the relationship between histological features and aging markers such as telomere length.

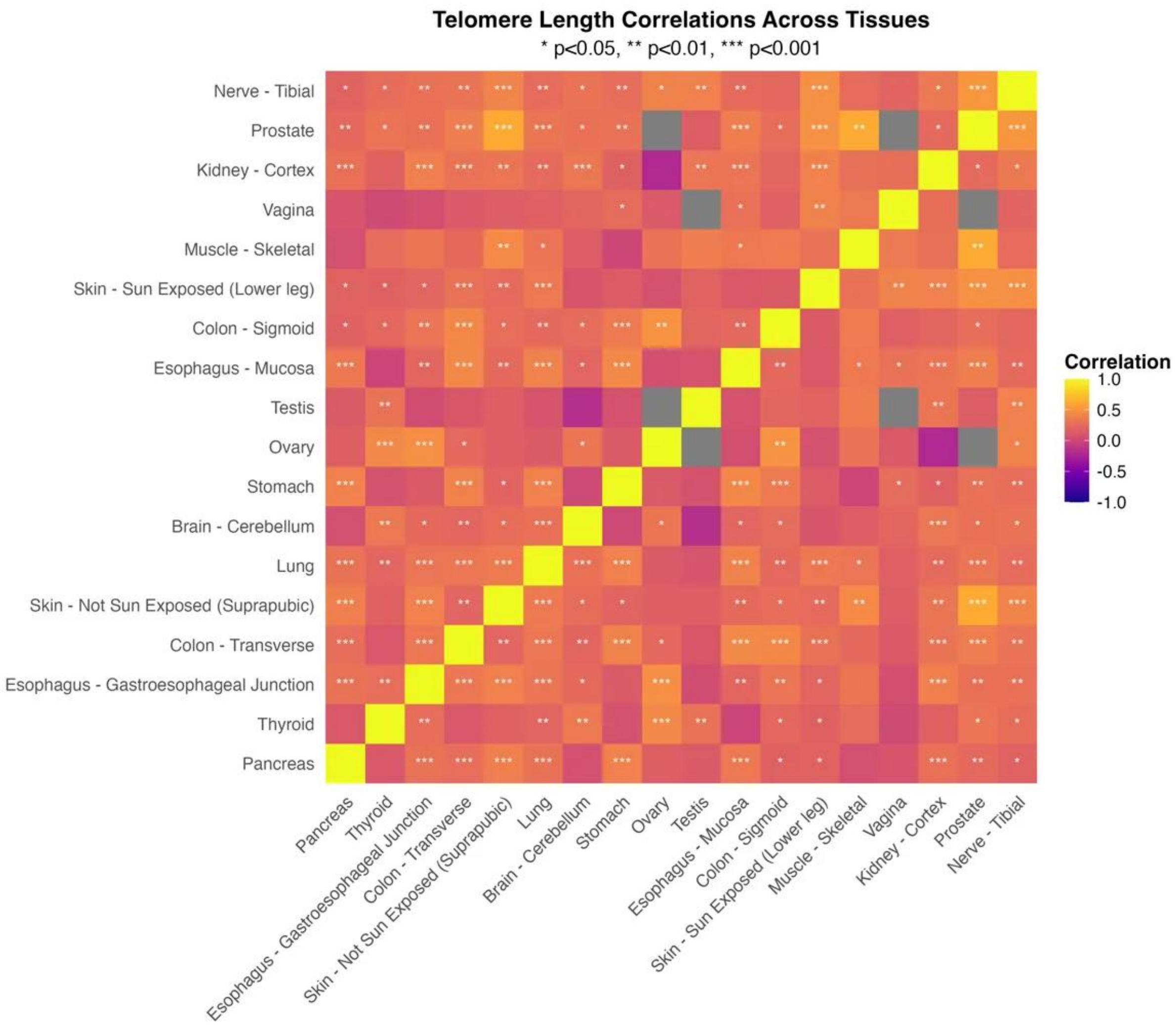


**Figure S3: Within-individual correlation analysis of telomere length across different tissues.** Heatmap displaying pairwise Pearson correlations of telomere length between different tissues collected from the same individuals in the GTEx dataset. The diagonal shows perfect self-correlation (r = 1.0), while off-diagonal elements represent how telomere lengths correlate between tissue pairs within individuals. Colors range from yellow (strong positive correlation, r = 1.0) to purple (strong negative correlation, r = -1.0), with gray squares indicating insufficient paired samples for correlation calculation. Statistical significance is indicated by asterisks (* $p<0.05$, ** $p<0.01$, *** $p<0.001$). The predominantly moderate positive correlations suggest systemic influence on telomere length maintenance, while the varying levels of significance across tissue pairs indicate tissue-specific regulation even within the same individual. This analysis spans 18 tissue types, with many tissue pairs showing statistically significant correlations.

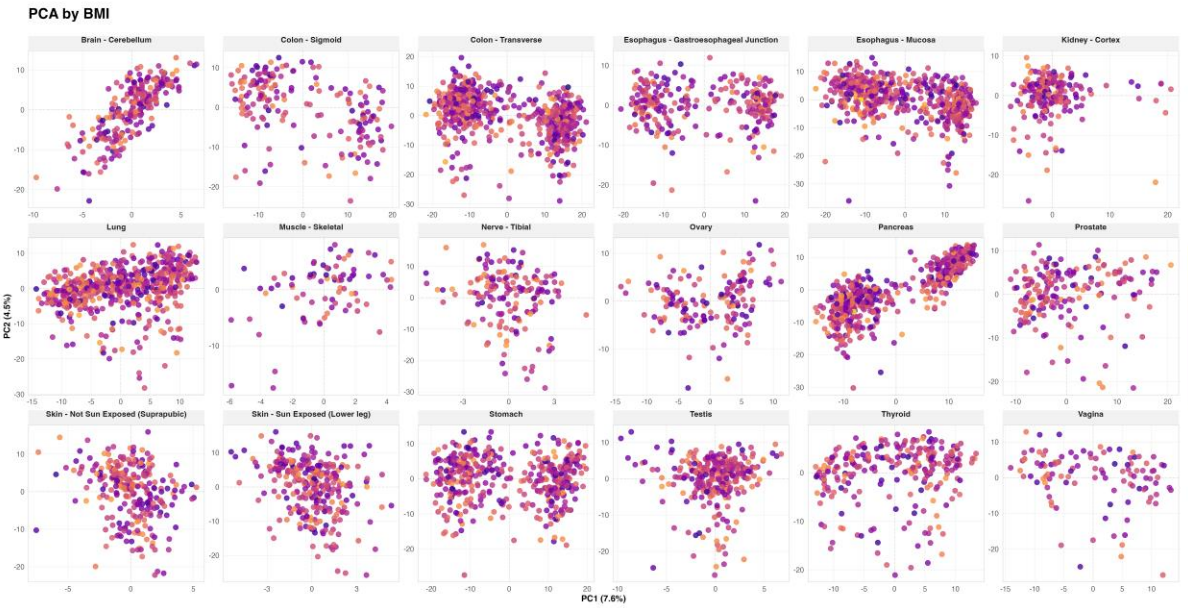


**Figure S4: Principal Component Analysis (PCA) of H&E image features shows limited association with BMI across tissues.** Visualization of the first two principal components (PC1 vs PC2) derived from UNI-extracted image features across 18 different GTEx tissues. Individual points represent distinct samples, with color indicating Body Mass Index (BMI) values (scale from purple [lower BMI] to yellow [higher BMI]). The relatively uniform distribution of BMI values across the PCA space in most tissues suggests that the primary sources of morphological variation captured by these components are not strongly associated with BMI.

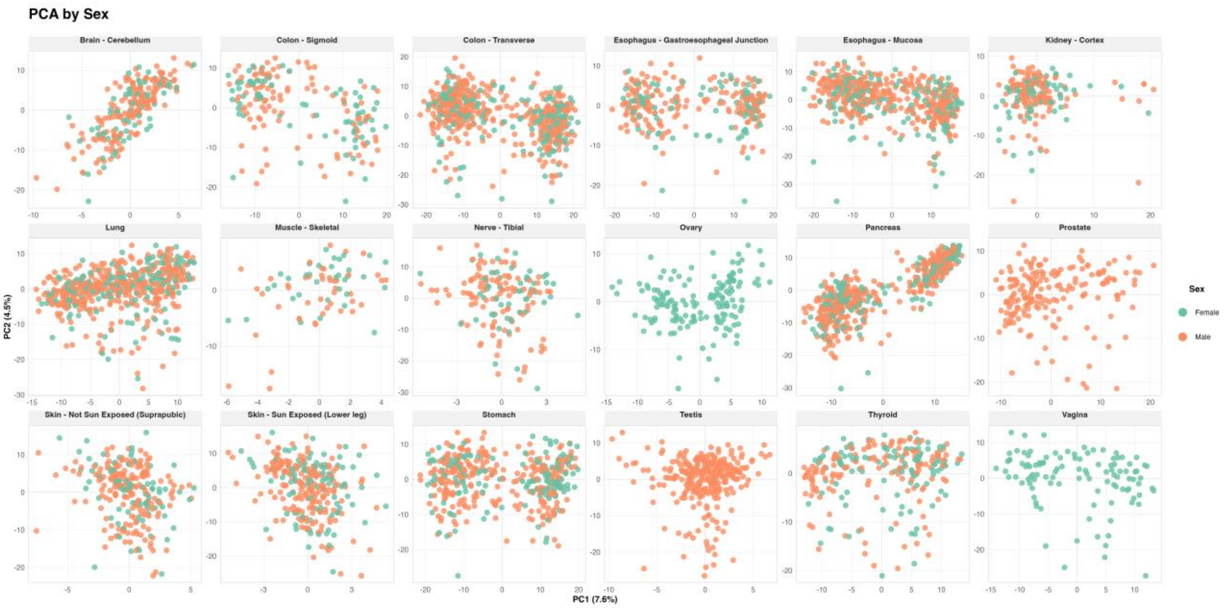


**Figure S5: Principal Component Analysis (PCA) of H&E image features shows no clear sex-based patterns across tissues.** Visualization of the first two principal components (PC1 vs PC2) derived from UNI-extracted image features across 18 different GTEx tissues. Individual points represent distinct samples, colored by sex (blue: male, red: female). The relatively uniform distribution of male and female samples across the PCA space in all tissues suggests that the primary sources of morphological variation captured by these components are not strongly associated with biological sex.

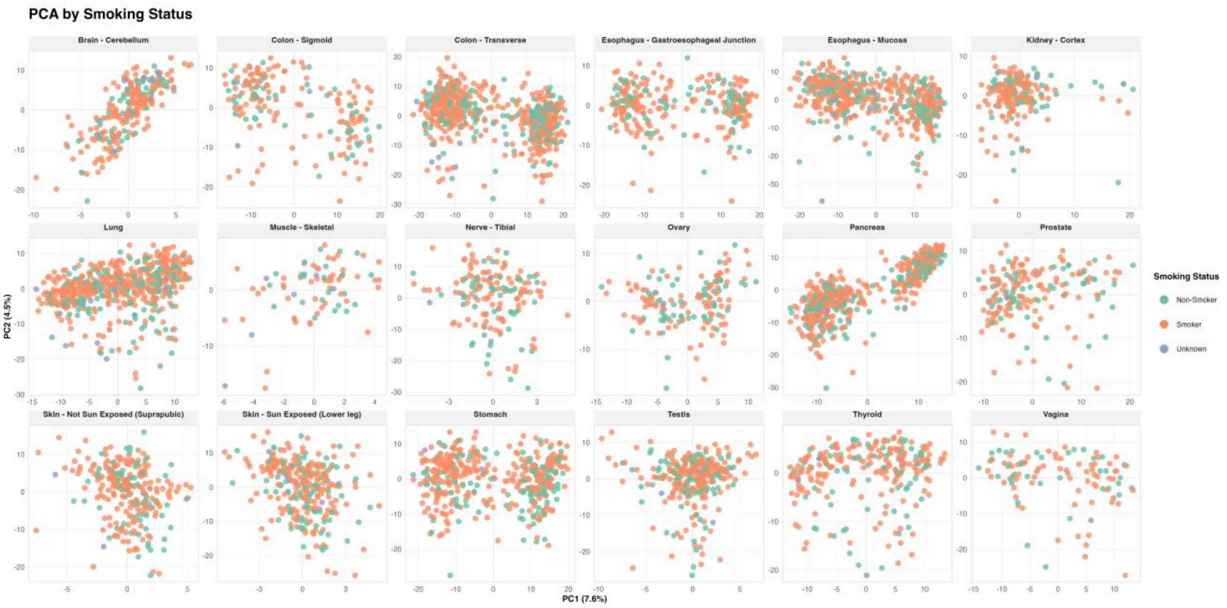


**Figure S6: Principal Component Analysis (PCA) of H&E image features shows no clear association with smoking status across tissues**. Visualization of the first two principal components (PC1 vs PC2) derived from UNI-extracted image features across 18 different GTEx tissues. Individual points represent distinct samples, colored by smoking status (light green for non-smoker, light orange for smoker and light blue for unknown). The distribution of smoking status categories appears random across the PCA space in all tissues, suggesting that the primary morphological variations captured by these components are not strongly influenced by smoking history.

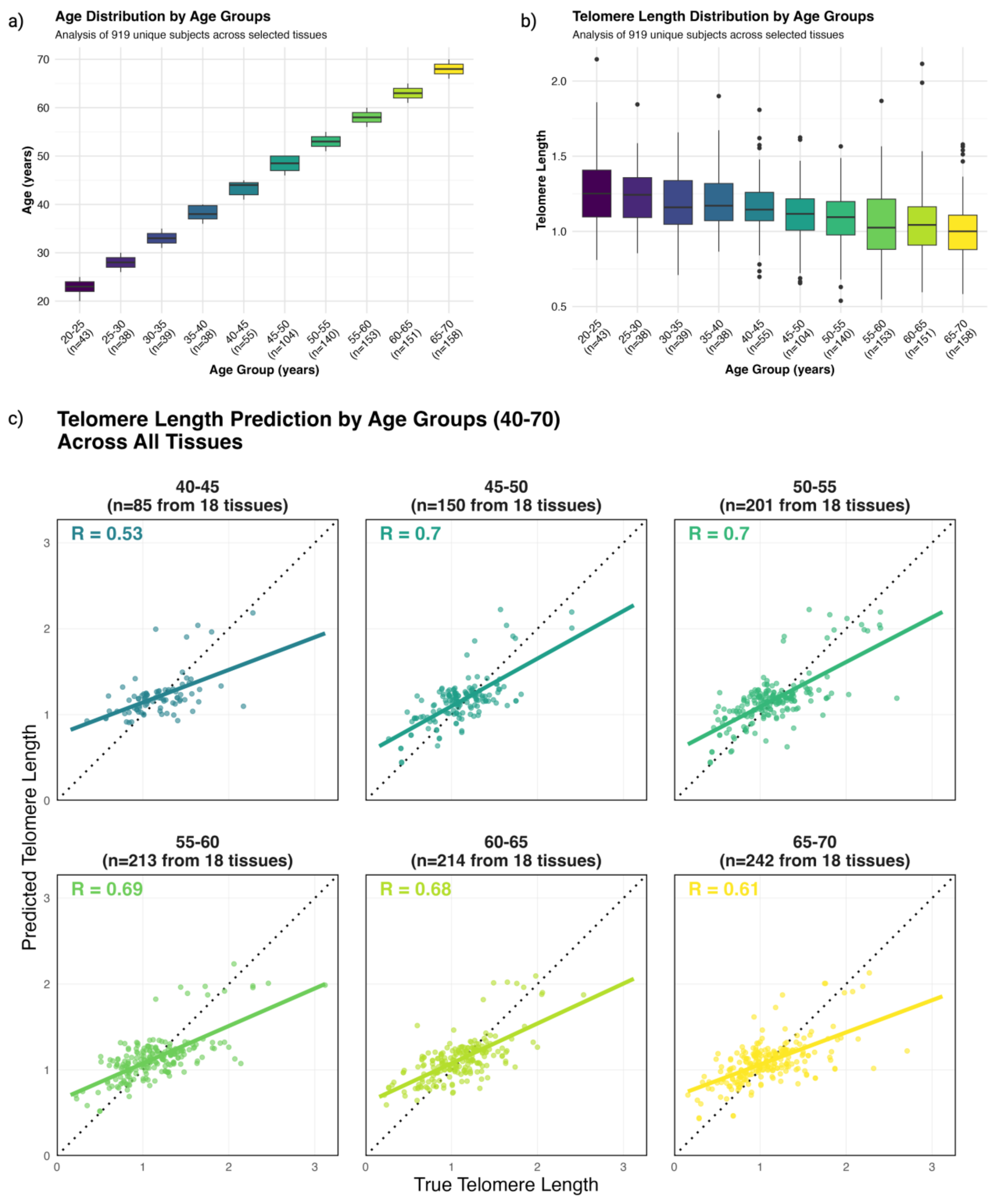


**Figure S7: Analysis of age distribution and telomere length predictions across different age groups.** The figure consists of three components: (**a**) A box plot showing the age distribution within each 5-year age group from 20-70 years, demonstrating the even stratification of the study population. (**b**) A box plot illustrating the distribution of telomere lengths across age groups, with an overall correlation coefficient of -0.229, suggesting a weak negative correlation between age and telomere length. (**c**) Six scatter plots showing the relationship between predicted and true telomere length (presented as Z-scores) for age groups between 40-70 years. Each subplot includes the Pearson correlation coefficient (r) and sample size (n). The correlation strength varies across age groups, with the highest correlation observed in the 45-50 age group (r = 0.77, n = 15) and the lowest in the 40-45 age group (r = -0.36, n = 5). The diagonal dashed line represents perfect prediction. These results demonstrate that prediction accuracy varies across age groups, with most groups showing moderate to strong positive correlations between predicted and actual telomere lengths.

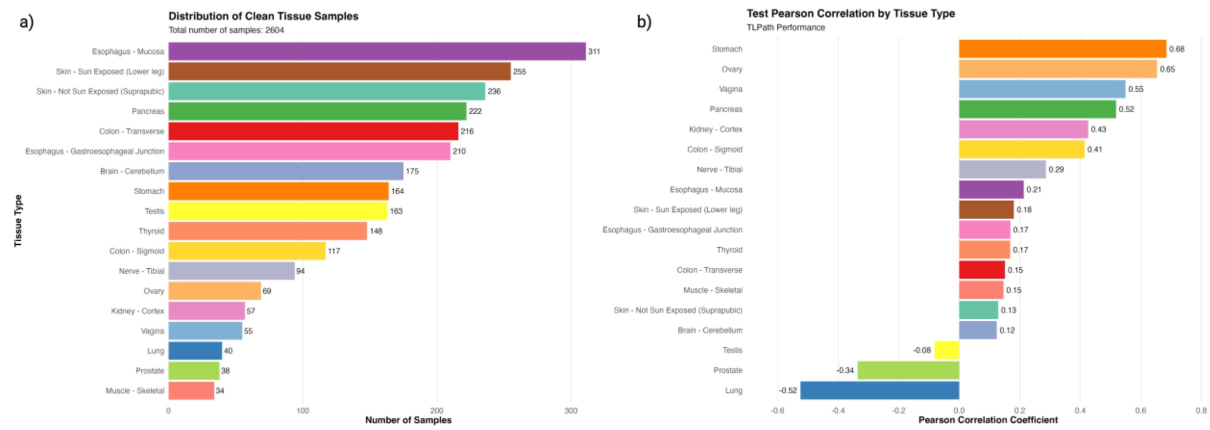


**Figure S8: Performance of TLPath using clean tissue samples.** (**a**) The distribution of samples for each tissue is shown. Filtering was done using each sample's pathology notes, excluding samples with pathology notations of autolysis and disease-related terminology. The total number of samples after this filtering was 2,604. (**b**) TLPath's performance is shown using the cleaned dataset. The top four tissues include stomach (r = 0.68), ovary, (r = 0.65), vagina (r = 0.55) and pancreas (r = 0.52). Tissues with no predictive ability in this dataset (r < 0) include testis (r = -0.06), prostate (r = -0.34) and lung (r = -0.52).

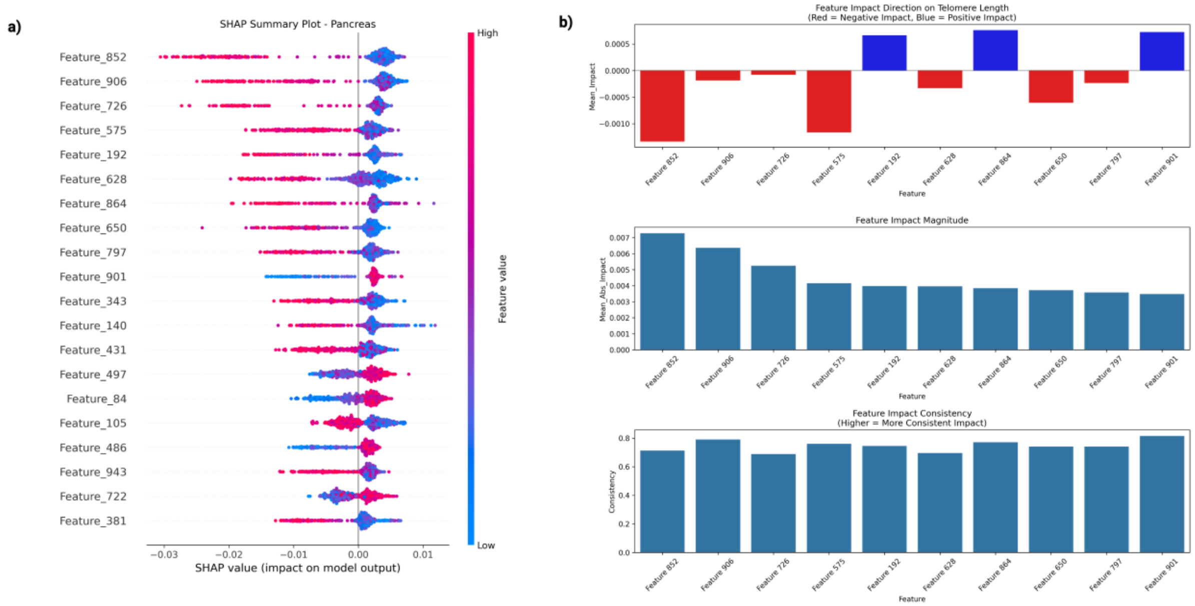


**Figure S9: Feature impact analysis on telomere length predictions in pancreas tissue**. (**a**) Distribution of SHAP values across samples for the same top 20 features, ordered by mean absolute SHAP value. Each point represents a sample, with position indicating the SHAP value's magnitude and direction (negative values decrease predictions, positive values increase them). Point color shows the original feature value (blue = low, red = high). This visualization reveals how feature effects vary across samples and whether the impact is consistent across different feature values. For example, high values of feature 906 (red points) consistently contribute to decreased telomere length predictions (negative SHAP values). (**b**) Three plots showing the feature impact direction, feature impact magnitude, and feature impact consistency of the top 10 features. Feature impact direction is the mean impact of all SHAP values for each corresponding feature. Feature impact magnitude is the absolute value of all SHAP values for each corresponding feature. Feature impact consistency is measured by analyzing how stable a feature's SHAP values remain across different samples and cross-validation folds, considering both the magnitude and direction of its effect on telomere length predictions.

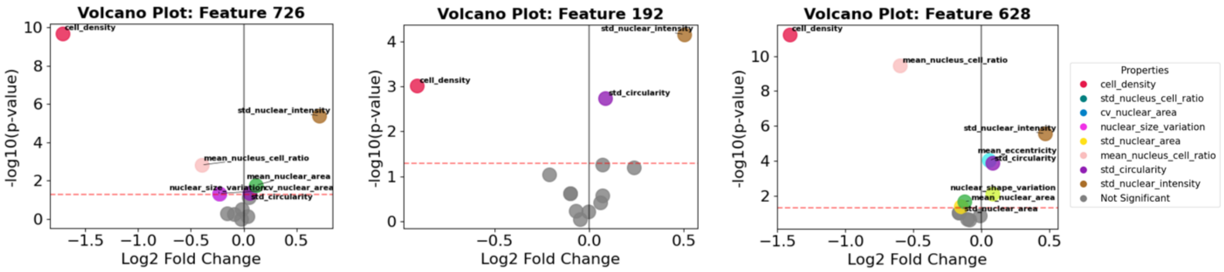


**Figure S10: QuPath-based interpretability of TLPath's top features.** From both long and short telomere samples, we extracted the top 5 patches per feature from each tissue sample. The plot compares the long versus short telomere patches for features 726, 192 and 628, depicting log fold changes (x-axis) against statistical significance (-log10 p-value, y-axis). Points right of zero indicate properties elevated in long telomere samples, while points left of zero show properties enriched in short telomere samples. The plots highlight differential cellular morphology characteristics between long and short telomere conditions.

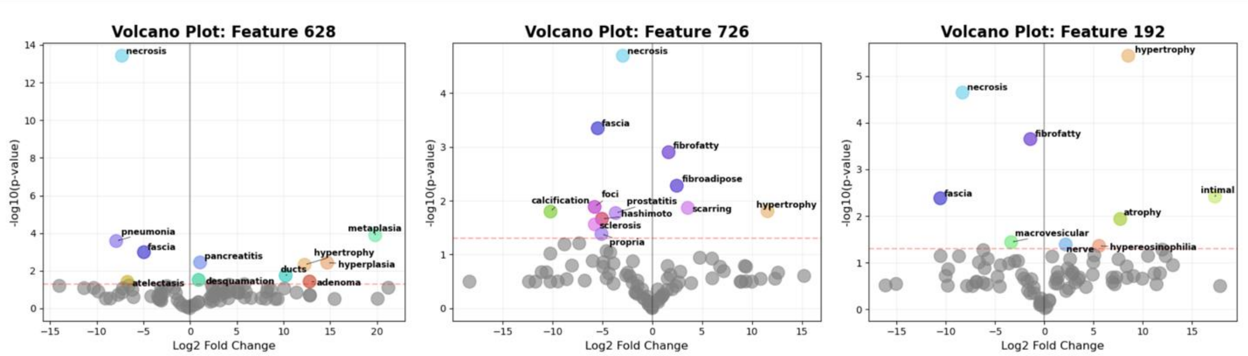


**Figure S11: CONCH word analysis volcano plot.** From both long and short telomere samples, we extracted the top 5 patches per feature from each tissue sample. The patches along with 86 histological text terms are given to the model CONCH 1.5 and the average confidence score is calculated for all terms, across all patches. The plots highlight the association of our terms from GTEx pathology reports to features patches 726, 192 and 628, by with long and short telomere from WSI. Points right of zero indicate elevated association in long telomere samples, while points left of zero indicate an elevation in short telomere samples.

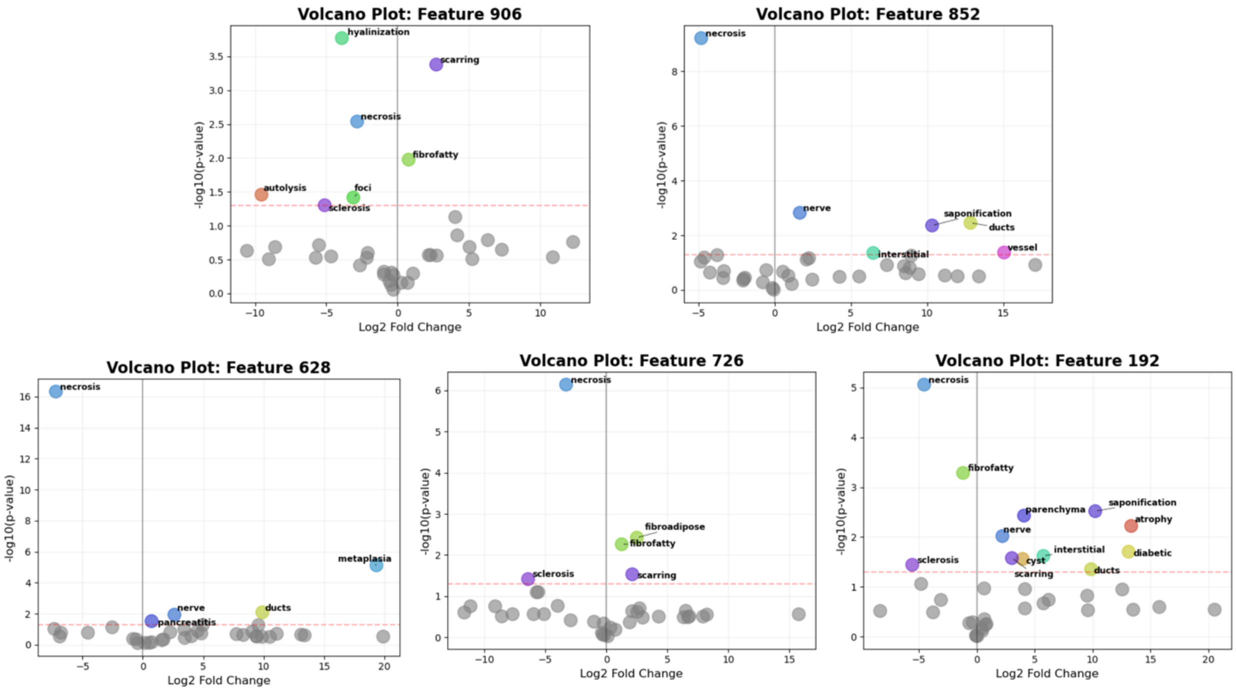


**Figure S12: CONCH word analysis for pancreas specific terms.** From both long and short telomere samples, we extracted the top 5 patches per feature from each tissue sample. The patches along with 37 pancreas specific histological text terms are given to the model CONCH 1.5 and the average confidence score is calculated for all terms, across all patches. The plots highlight the association of our pancreas specific terms from GTEx pathology reports to feature patches 906, 852, 726, 192 and 628, by with long and short telomere from WSI. Points right of zero indicate elevated association in long telomere samples, while points left of zero indicate an elevation in short telomere samples. We gather our pancreas specific terms as descried in 4.6.2, but now, only consider terms in pathology notes and categories under pancreas tissues. We note that the morphological feature necrosis is higher in samples with short telomere lengths. This is consistent with our previous analysis. Terms features ducts and nerve are observed in three patch features (192, 628, 852) to be higher in samples with long telomere lengths, whereas sclerosis (features 192, 762, 906), is higher in samples with short telomere length.

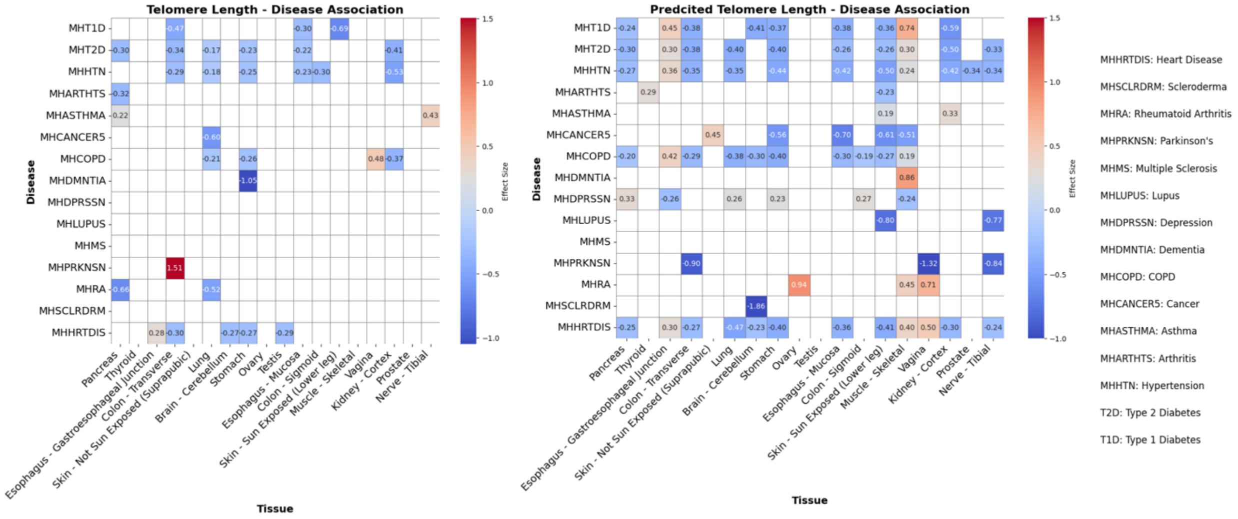


**Figure S13: Disease associations with tissue-specific telomere lengths.** We investigated the relationship between tissue-specific telomere length and various diseases using two approaches: measured telomere length and model-predicted telomere length. The associations are quantified using Cohen's d effect size, where positive values (shown in red) indicate longer telomere length in disease cases compared to controls, and negative values (shown in blue) indicate shorter telomere length in disease cases. **(a)** The analysis revealed several significant tissue-specific associations. Most notably, Parkinson's disease demonstrated the strongest positive association with pancreatic tissue telomere length (d = 1.51), suggesting substantially longer telomeres in pancreatic tissue of Parkinson's patients. Type 1 Diabetes exhibited consistent negative associations across multiple tissues, including thyroid (d = -0.47), brain-cerebellum (d = -0.30), and ovary (d = -0.69). COPD showed negative associations with brain-related tissues, specifically in brain-cerebellum (d = -0.21) and brain-cortex (d = -0.26), while displaying a positive association with skeletal tissue (d = 0.48). Heart Disease demonstrated a pattern of negative associations in brain tissues (d = -0.27) with a positive association in thyroid tissue (d = 0.28). **(b)** The second heatmap presents the association between predicted telomere length, derived from TLPath, and disease associations across tissues. Parkinson's disease exhibited strong negative associations in skeletal tissue (d = -1.32) and nerve-tibial (d = -0.84), while showing a positive association in gastroesophageal junction (d = 0.90). Scleroderma demonstrated a strong positive association with brain-cerebellum (d = 1.86). Lupus displayed significant positive associations in muscle-skeletal (d = 0.80) and nerve-tibial (d = 0.77) tissues. Cancer showed consistent positive associations across multiple tissues, including ovary (d = 0.70), esophagus-mucosa (d = 0.61), and skeletal (d = 0.51). Both Type 1 and Type 2 Diabetes exhibited widespread negative associations across various tissues, with effect sizes ranging from -0.24 to -0.50.

| Property Name | Description |
|---|---|
| **cell_count** | Total number of cells detected in the patch |
| **cell_density** | Number of cells per unit area |

| | |
|---|---|
| **mean_nuclear_area** | Average area of nuclei in the patch |
| **std_nuclear_area** | Standard deviation of nuclear areas in the patch |
| **cv_nuclear_area** | Coefficient of variation of nuclear areas |
| **mean_circularity** | Average circularity of nuclei |
| **std_circularity** | Standard deviation of nuclear circularity |
| **mean_eccentricity** | Average eccentricity (elongation) of nuclei |
| **nuclear_size_variation** | Variation in nuclear sizes in a patch |
| **nuclear_shape_variation** | Variation in nuclear shapes in a patch |
| **mean_nucleus_cell_ratio** | Average ratio of nucleus size to cell size |
| **std_nucleus_cell_ratio** | Standard deviation of nucleus-to-cell ratio in the patch |
| **mean_nuclear_intensity** | Average intensity of nuclear staining in the patch |
| **std_nuclear_intensity** | Standard deviation of nuclear staining intensity in the patch |

**Table S1: Quantitative metrics for nuclear and cellular analysis per patch.** This table presents a comprehensive set of morphological and organizational metrics extracted from whole slide images using QuPath cell detection. The metrics encompass multiple aspects of cellular and nuclear characteristics, providing a quantitative framework for tissue architecture analysis. Each row in the table contains three columns: (1) the property name as implemented in the code, (2) a description of what the property represents biologically or technically.
